# Paternal diet shapes paternal and maternal transcriptomes in the early embryo through parallel sperm-associated mechanisms

**DOI:** 10.64898/2026.09.23.753702

**Authors:** Leonard C Steg, Inés Blanc Giró, Kerem Uzel, Rodrigo G Arzate-Mejia, Isabelle M Mansuy

## Abstract

Life experiences in fathers can influence phenotypes in the offspring. While it is known that acquired paternal information can be transferred to descendants via the germline, how such transfer operates between sperm and embryo remains unclear. We examined the effects of low protein diet (LPD) on males of different genotypes and their hybrid offspring generated by reciprocal outcross and on their genome. We found that LPD induces genotype-specific effects in fathers and different phenotypes in their male and female offspring. Allele-specific analyses revealed that paternal LPD modifies paternal and maternal transcriptomes differently in early two-cell embryos. Changes in the paternal transcriptome overlap with the sperm transcriptome and are associated with sperm chromatin features, suggesting that sperm-inherited transcripts and chromatin states may shape the paternal transcriptome in the embryo. The maternal transcriptome, by contrast, is linked to sperm-derived small RNAs, consistent with a possible role of sperm-derived small RNAs in modulating maternal transcript levels in the embryo. Analyses of undifferentiated spermatogonia populations suggest that LPD-induced sperm states can originate from early stages of germ cells. These findings provide evidence that paternal experiences can influence the early embryo via processes involving each parental allele differently.

**Graphical abstract:** 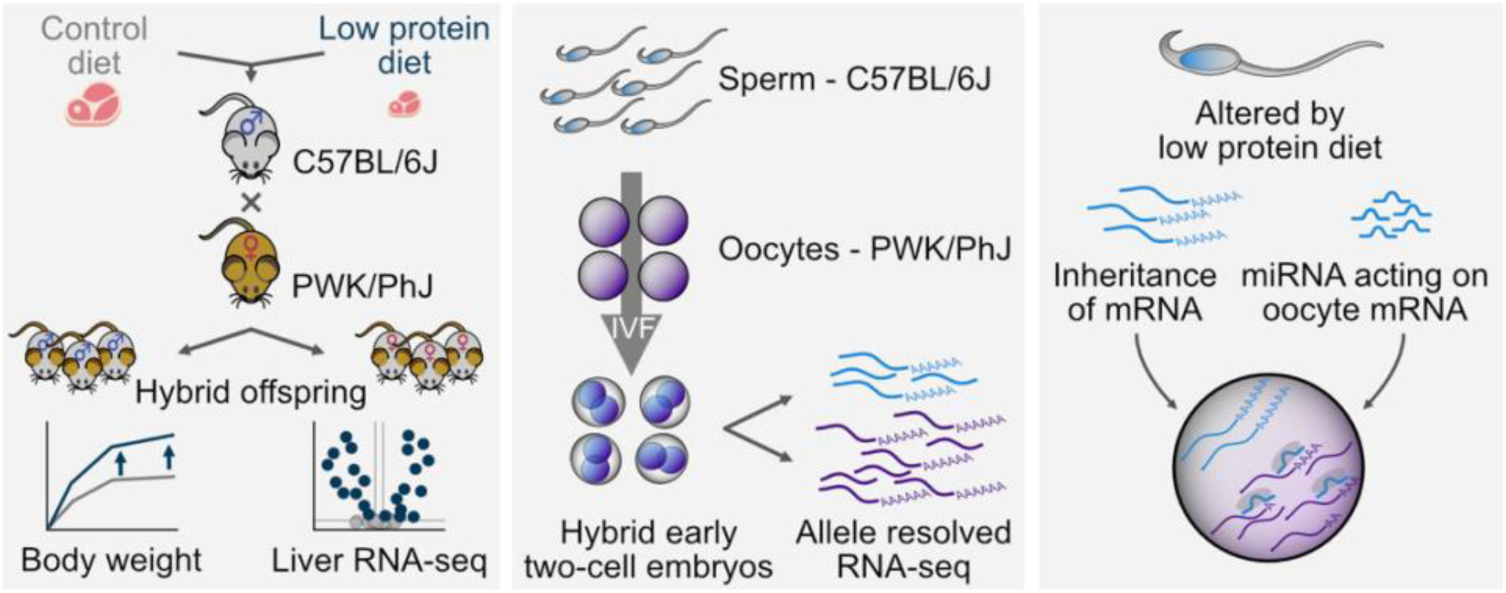

## Introduction

Life conditions and environmental factors experienced by parents influence their own physiology and health and can also impact their offspring. Whether adverse or beneficial, they can modulate disease susceptibility in parents and offspring ^1–3^. In rodent models, paternal exposure to poor diet ^4,5^, stress ^6,7^, endocrine disruptors ^8,9^ or antibiotics ^10,11^ has been shown to modify offspring phenotypes, sometimes across several generations ^12^. Such effects can involve the transfer of molecular information from father to offspring through the germline, independent of changes in the DNA sequence. In rodents and humans, exposure has been shown to modify the sperm transcriptome and epigenome including RNA ^13–15^, DNA methylation (DNAme) ^7,16–18^, histone post-translational modifications (PTMs) ^18,19^ and three-dimensional chromatin configuration ^20^. Such molecular changes are thought to contribute to the transfer of signals from father to its direct descendants. Among them, RNA and specific histone PTMs have been linked with effects on gene regulation in embryos ^13,19,21^. However, whether paternal signals induced in sperm can be transferred to the offspring remains unclear.

Shortly after fertilization, the early embryo is largely transcriptionally inactive and its RNA load is composed of transcripts inherited from the maternal and paternal gametes. This parental contribution is unequal and oocyte-derived transcripts are far more abundant than those coming from sperm ^22^. At this early stage of development prior to zygotic genome activation (ZGA), the paternal genome undergoes extensive chromatin remodeling then becomes transcriptionally active ^22,23^. In mice, ZGA occurs in two waves, with a minor wave at the one-cell stage followed by a major wave at the late two-cell stage that marks the onset of full embryonic transcriptional activity. Epigenetic states at maternal and paternal alleles markedly differ during early embryonic development with differences in chromatin compaction, non-canonical distributions of histone PTMs and contrasting DNAme patterns ^23^.

To precisely determine how paternal exposure influences the early embryo, it is necessary to distinguish the paternal and maternal contribution to the embryo transcriptome and not treat it as a single unit. Past studies have examined allele-specific effects of exposure after ZGA at the four-cell stage or post-implantation ^24,25^, which is too late to identify the specific paternal contribution. We used a reciprocal outcross strategy between parents from two genetically divergent mouse strains, C57BL/6J and PWK/PhJ, to generate hybrid embryos and analyze the effects of paternal low protein diet (LPD) before major ZGA. LPD is a condition that reliably alters body weight, liver metabolism and cardiovascular functions in exposed males and their progeny in mice ^4,26–29^. Our results show that paternal LPD produces strain-dependent responses in exposed males but consistent intergenerational effects on body weight in the offspring from both strains. In early embryos, paternal diet induced distinct transcriptional changes on the paternal and maternal genomes that may originate in germ cell precursors.

## Results

### Strain-specific paternal responses and shared sex-specific intergenerational effects of LPD

To enable allele-specific analyses in embryos, we generated hybrid offspring using C57BL/6J or PWK/PhJ males reciprocally bred with PWK/PhJ or C57BL/6J females, respectively. At weaning (3-week), C57BL/6J and PWK/PhJ males were fed LPD (9% protein) or control diet (18.5% protein) for 10 weeks and at 3-months of age, the males were mated with naïve adult females of the respective other strain fed control diet (Fig. 1, A and B). We observed that LPD decreases body weight in C57BL/6J males (Fig. 1A) as shown before ^4,30^ but has no effect in PWK/PhJ males (Fig. 1B). In the hybrid offspring from C57BL/6J males fed LPD crossed with naïve PWK/PhJ females, males gained more weight than controls across time (Fig. 1C) but females did not (Fig. 1D), as previously reported ^28^. Similar effects were observed in the progeny of PWK/PhJ males fed LPD crossed with naïve C57BL/6J females, with males but not females gaining more weight than controls (Fig. 1E, F). At 3-months of age, the offspring of C57BL/6J or PWK/PhJ LPD-fed males were respectively 8.9% and 6.2% heavier than the corresponding control offspring (Fig. 1C, E).

**Figure 1.**
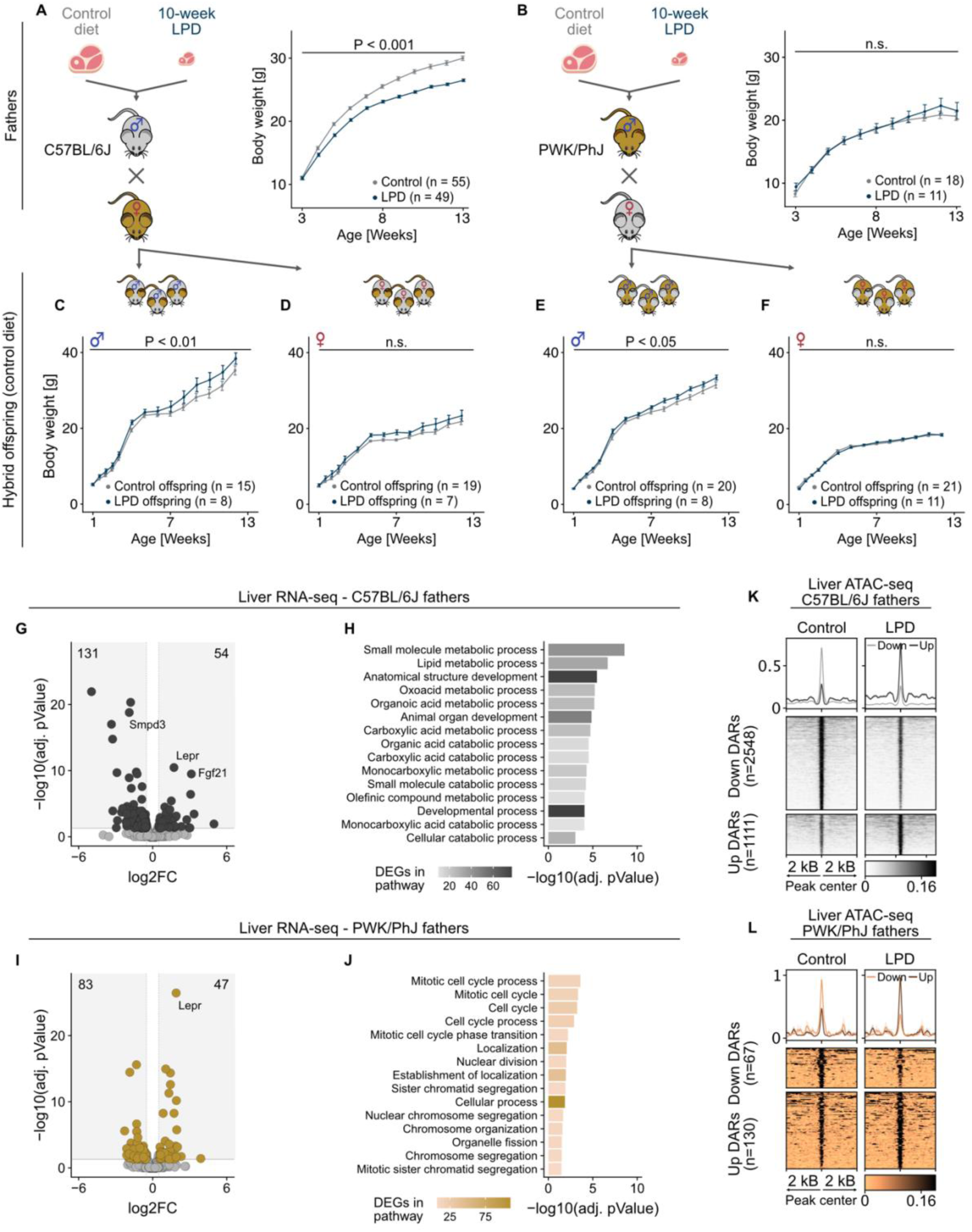
Effects of LPD on body weight in fathers and their hybrid offspring and on liver transcriptome and chromatin accessibility in fathers. (**A** and **B**) Experimental design and body weight trajectories of (A) C57BL/6J and (B) PWK/PhJ fathers fed a control diet or 10-week LPD prior to mating. Data are presented as mean ± SEM. Statistical analyses include a mixed-effects model (*Bodyweight* ∼ Group:Age + Age + (1|Animal) + (1|Litter)), *P* values were extracted from Group:Age interaction, representing changes in body weight during diet exposure. (C–F) Body weight trajectories of hybrid offspring derived from control or LPD-fed fathers. (C) Male and (D) female offspring of C57BL/6J fathers. (E) Male and (F) female offspring of PWK/PhJ fathers. Data are presented as mean ± SEM. Statistical analyses include a mixed-effects model (*Bodyweight* ∼ Group:Age + Age + (1|Animal) + (1|Father)), *P* values correspond to Group:Age interaction, representing changes in body weight during development. (G) Volcano plot of DEGs (adjusted *P* < 0.05 and abs(log2FC) > 0.5) in liver of C57BL/6J fathers fed LPD compared to control diet. Numbers indicate the count of down- and upregulated DEGs. Control: n = 5, LPD: n = 5. (H) Top 15 biological process GO terms most significantly enriched among DEGs in C57BL/6J fathers. (I) Volcano plot of DEGs (adjusted *P* < 0.05 and abs(log2FC) > 0.5) in liver of PWK/PhJ fathers fed LPD compared to control diet. Numbers indicate the count of down- and upregulated DEGs. Control: n = 5, LPD: n = 4. (J) Top 15 biological process GO terms most significantly enriched among DEGs in PWK/PhJ fathers. (K and L) Heatmaps showing normalized ATAC-seq signal across DARs (adjusted *P* < 0.05 and abs(log2FC) > 1) between control and LPD groups in (K) C57BL/6J (Control: n = 5, LPD: n = 5) and (L) PWK/PhJ (Control: n = 4, LPD: n = 4) fathers. Each row represents a 4-kb region centered on the DAR midpoint (±2 kb), ordered by mean ATAC-seq signal.

Liver is an important metabolic organ that dynamically responds to dietary intake ^31^. We profiled the transcriptome of liver from C57BL/6J males fed LPD and detected 185 differentially expressed genes (DEGs) (adjusted P < 0.05, abs(log2FC) > 0.5) with 131 being downregulated and 54 upregulated (Fig. 1G). Gene ontology (GO) enrichment analysis showed that DEGs are significantly enriched for metabolic and catabolic processes and for developmental pathways (Fig. 1H). Among downregulated genes was *Smpd3*, which encodes an enzyme involved in lipid homeostasis that responds to dietary perturbations (Fig. S1A) ^32^. *Fgf21,* a metabolic regulator that can be induced by protein restriction and promotes weight loss was upregulated, consistent with the decreased weight in LPD-fed C57BL/6J males (Fig. S1A) ^33^. In PWK/PhJ males fed LPD, 130 DEGs were detected (adjusted P < 0.05, abs(log2FC) > 0.5) with 83 being downregulated and 47 upregulated (Fig. 1I). DEGs were primarily enriched in cell cycle and mitosis-related processes (Fig. 1J). Among the upregulated DEGs was *Lepr*, which encodes a leptin receptor responsive to protein restriction ^34^ (Fig. S1B). Notably, *Lepr* was one of the few genes upregulated in both C57BL/6J and PWK/PhJ males (Fig. S1A), but overall DEGs minimally overlapped between the two strains (5 downregulated and 2 upregulated; Fig. S1C), suggesting distinct transcriptional response to the same dietary challenge.

Gene transcription is tightly coupled to chromatin organization, and open chromatin at regulatory elements is a prerequisite for transcription factor (TF) binding to activate or repress gene expression^35^. Thus, we profiled chromatin accessibility in liver using assay for transposase-accessible chromatin (ATAC)-seq. In C57BL/6J males, 2,548 genomic regions had decreased accessibility (adjusted P < 0.05, log2FC < –1) and 1,111 regions had increased accessibility (adjusted P < 0.05, log2FC > 1) (Fig. 1K). Differentially accessible regions (DARs) were predominantly located in intronic and distal intergenic regions, with only a small fraction corresponding to promoters, suggesting that distal regulatory elements are primarily affected by LPD (Fig. S2A). To further characterize these regions, we annotated DARs using publicly available PTMs datasets from mouse liver (Fig. S2B). Both closing and opening DARs were marked by H3K4me1 and depleted of H3K4me3, a histone PTM combination typical of enhancer regions ^36^. About half of opening DARs also carried H3K27ac, suggesting that these regions are active enhancers ^36^. In liver from PWK/PhJ males, only 197 DARs were detected, 67 with decreased accessibility and 130 with increased accessibility (Fig. 1L). These DARs were as well enriched in intronic and distal intergenic regions (Fig. S2C) and most were marked by H3K4me1 with a subset also carrying H3K27ac, also pointing to active enhancers (Fig. S2D). We next examined the enrichment of TF binding motifs within DARs of both strains. TFs were selected for their established role in hepatic metabolism^37,38^ or their implication in LPD effects ^29^. In C57BL/6J males, motifs for all assessed TFs including peroxisome proliferator-activated receptors (PPARα and PPARγ) and multiple hepatic nuclear factors (HNFs) were enriched in both opening and closing DARs (Fig. S2E). In contrast in PWK/PhJ males, only PPARα and HNF4A motifs were enriched in both closing and opening DARs. Other TFs motifs (PPARγ, FOXA1, HNF6, HNF6b and activating TF7 (ATF7)) were enriched in either closing or opening DARs, and HNF1 and HNF1B motifs were not enriched in any DAR (Fig. S2E). Given that C57BL/6J males and their offspring show the strongest physiological and molecular effects of LPD, this strain was used for further analyses.

### Paternal LPD induces sex-specific transcriptional and chromatin changes in offspring liver

To better characterize the intergenerational effects of paternal LPD, we next profiled the transcriptome and chromatin accessibility of liver from adult hybrid offspring from C57BL/6J fathers fed LPD crossed with PWK/PhJ mothers fed control diet. In male offspring, paternal LPD altered the expression of 51 genes (adjusted P < 0.05, abs(log2FC) > 0.5), with 13 being downregulated and 38 upregulated (Fig. 2A). DEGs were enriched for metabolic pathways, particularly lipid metabolic process and other lipid-related processes (Fig. 2B) known to be altered by paternal LPD ^4,26,29^. The gene with the most significant upregulation was *Pnpla3*, previously found to be upregulated in offspring liver after paternal LPD ^29^. *Pnpla3* has been associated with obesity in mice ^39^ and its human ortholog *PNPLA3* with liver fat accumulation exacerbated by weight gain ^40^. The gene with the largest fold change was *Cyp2b9*, implicated in body weight gain after high fat diet in mice ^41^ (Fig. S3A). These transcriptional changes align with the increased body weight observed in male offspring of fathers fed LPD (Fig. 1C). Notably, only a subset of individual DEGs identified in male offspring overlapped with DEGs observed in their C57BL/6J fathers (3 downregulated and 4 upregulated; Fig. S3B), which is not surprising given their opposite body weight phenotype. In female offspring, 41 DEGs were detected, 21 downregulated and 20 upregulated genes (Fig. 2C), and were enriched for metabolic pathways related to lipids (Fig. 2D). There was no overlap between DEGs in male and female offspring (Fig. S3D).

**Figure 2.**
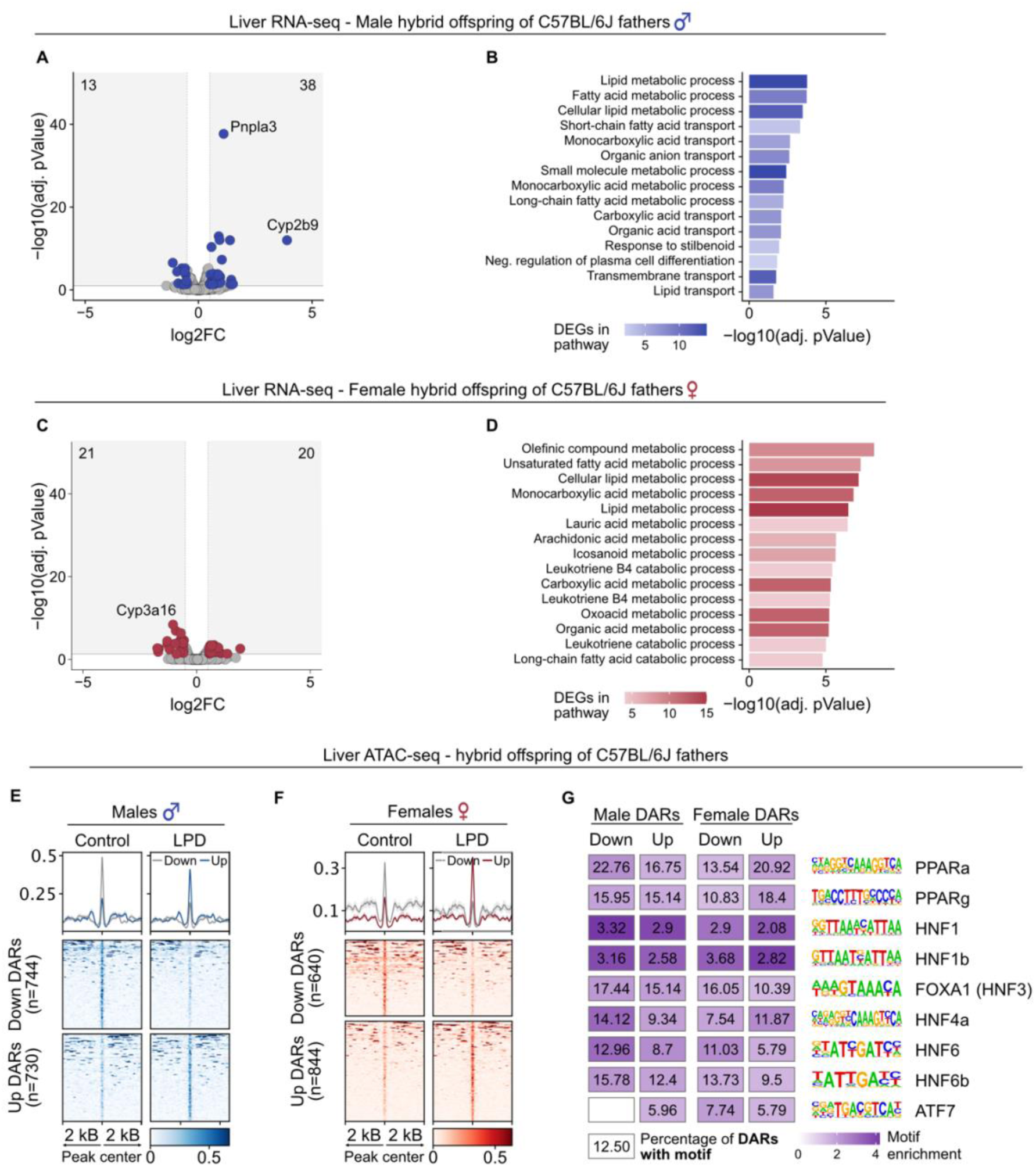
Analyses of liver transcriptome and chromatin accessibility in offspring of LPD-fed C57BL/6J fathers. (**A** and **B**) Effects of paternal LPD on the liver transcriptome of male offspring. Control: n = 10, LPD: n = 7. (**A**) Volcano plot of DEGs (adjusted *P* < 0.05 and abs(log2FC) > 0.5) in liver of male offspring of C57BL/6J fathers fed LPD compared to control diet. Numbers indicate the count of down- and upregulated DEGs. (**B**) Top 15 biological process GO terms most significantly enriched among DEGs in male offspring. (**C** and **D**) Effects of paternal LPD on the liver transcriptome of female offspring. Control: n = 10, LPD: n = 6. (**C**) Volcano plot of DEGs (adjusted *P* < 0.05 and abs(log2FC) > 0.5) in liver of female offspring of C57BL/6J fathers fed LPD compared to control diet. Numbers indicate the count of down- and upregulated DEGs. (**D**) Top 15 biological process GO terms most significantly enriched among DEGs in female offspring. (**E** and **F**) Heatmaps showing normalized ATAC-seq signal across DARs (*P* < 0.05 and abs(log2FC) > 1) between control and LPD groups in (**E**) male (Control: n = 10, LPD: n = 8) and (**F**) female (Control: n = 10, LPD: n = 6) offspring of C57BL/6J fathers fed LPD compared to control diet. Each row represents a 4-kb region centered on the DAR midpoint (±2 kb), ordered by mean ATAC-seq signal. (**G**) Enrichment of metabolic TF motif occurrence in DARs identified in male and female offspring of C57BL/6J fathers fed LPD compared to control diet. Only motifs with significant enrichment after multiple testing correction are shown. Color scale indicates fold enrichment. Numbers indicate the percentage of DARs containing the motif. Empty panels indicate no significant TF motif enrichment.

Allele-specific analyses of DEGs using the hybrid genetic background of the offspring showed that most DEGs in males have biallelic expression but several upregulated genes have either paternal or maternal allele expression (Fig. S3E). In female offspring, the majority of DEGs were expressed biallelically and only a few had paternal or maternal allelic expression (Fig. S3F). No change in the pattern of allelic expression of DEGs was observed in male and female offspring, suggesting that paternal LPD does not alter parent-of-origin mechanisms of gene regulation (Fig. S3, G and H). Similar differential gene expression analyses were also conducted in the offspring of PWK/PhJ males fed LPD or control diet mated with naïve C57BL/6J females fed control diet. In both sexes, we observed fewer DEGs in the offspring of PWK/PhJ fathers (Males: 45 genes; Females: 21 genes) compared to the offspring of C57BL/6J fathers (Males: 51 genes, Females: 41 genes). However, DEGs of the offspring of PWK/PhJ fathers were enriched in similar metabolic pathways (Fig. S4, A to D).

ATAC-seq analyses identified 744 regions with decreased accessibility and 730 regions with increased accessibility in liver from male offspring (P < 0.05, abs(log2FC) > 1; Fig. 2E). Given the small number of FDR-significant regions, we used regions meeting a nominal P < 0.05 threshold for downstream analyses to explore potential regulatory trends. Most identified DARs were annotated to intronic or distal intergenic regions, with only a small fraction located at promoters (Fig. S5A). Annotation with liver histone PTMs showed that DARs are marked by H3K4me1, with a small central dip characteristic of enhancer-associated chromatin with bound TFs (Fig. S5B) ^36,42^. A subset of DARs also carried H3K27ac, suggesting active enhancers. In female offspring, 640 regions had decreased and 844 regions had increased accessibility (P < 0.05, abs(log2FC) > 1; Fig. 2F). Annotation of these DARs showed a genomic distribution and profile of histone PTMs similar to those observed in males (Fig. S5C and D). Reciprocal analyses assessing ATAC-seq signal in males at DARs identified in females and *vice versa* showed no consistent differences in accessibility in either direction (Fig. S5E and F). This points to largely sex-specific changes in chromatin accessibility. Analysis of TF motif enrichment of DARs revealed significant enrichment for metabolic TFs including PPARα, PPARγ and HNFs. This suggests that altered transcriptional regulation contributes to responses to paternal LPD (Fig. 2G).

### Paternal LPD influences the transcriptome of paternal origin in early embryos

We examined the consequences of paternal LPD on early embryos before major ZGA. Hybrid early 2-cell embryos were generated by *in vitro* fertilization (IVF) using sperm from C57BL/6J males fed LPD or control diet to fertilize oocytes from naïve PWK/PhJ females, and their transcriptome was profiled by single-embryo RNA-seq. Correct staging of early two-cell embryos was confirmed by mapping RNA-seq results onto a publicly available reference dataset spanning mouse (CAST/EiJ × C57BL/6J) preimplantation development, from zygote to late blastocyst ^43^ (Fig. S6A). Differential expression analysis using single nucleotide polymorphisms (SNPs) between C57BL/6J and PWK/PhJ strains allowed us to separate paternal and maternal transcripts in early embryos. These analyses identified 172 downregulated and 169 upregulated genes (adjusted P < 0.05, abs(log2FC) > 0.5) in the paternal transcriptome of embryos derived from LPD-fed fathers (Fig. 3A). DEGs were significantly enriched in pathways related to metabolic processes and gene expression (Fig. 3B). Among the DEGs were *Dot1l*, a chromatin remodeler linked to histone modifications and transcriptional control ^44^, and *Ash1l*, involved in histone methylation and transcriptional activation ^45^. To identify regulatory factors linked to these transcriptional changes, we conducted TF target enrichment analyses on DEGs from the paternal transcriptome. Several TFs whose targets are enriched in the DEGs, including HNF1A and FOXA1 (also known as HNF3A), were identified (Fig. 3C). These TFs also showed motif enrichments in DARs identified in fathers fed LPD and their adult offspring (Fig. 2G and Fig. S2E). The analyses further identified CREM, RUNX2 and MYC, previously reported to have targets enriched in blastocysts transcriptome after paternal low-protein high-sugar diet ^46^.

**Figure 3.**
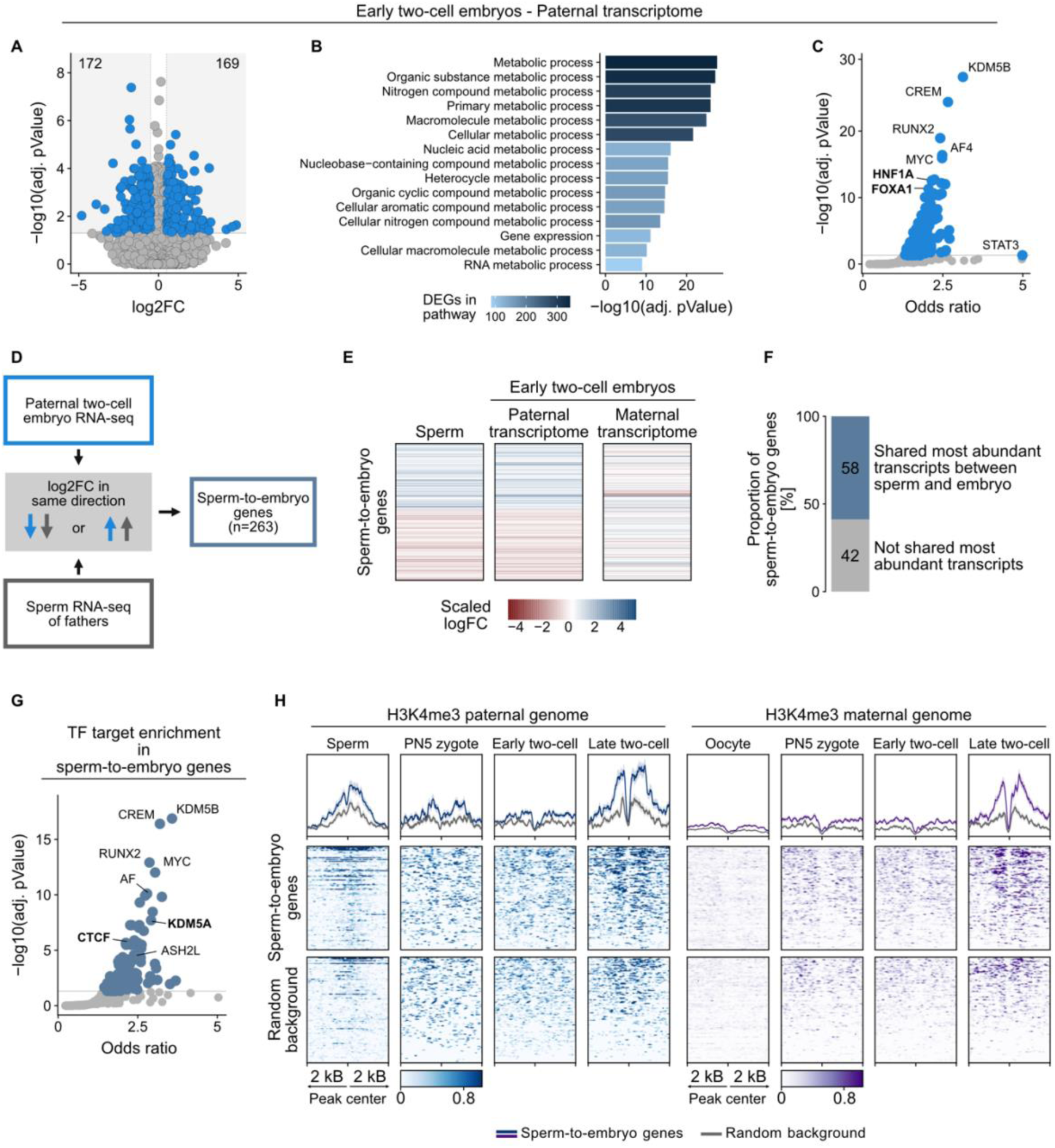
Paternal transcriptome analyses in early two-cell embryos and sperm from LPD-fed C57BL/6J fathers. (**A**) Volcano plot of DEGs (adjusted *P* < 0.05 and abs(log2FC) > 0.5) in the paternal transcriptome of early two-cell embryos derived from LPD-fed fathers compared to controls. Numbers indicate the count of down- and upregulated DEGs. Control: n = 42, LPD: n = 45. (**B)** Top 15 biological process GO terms most significantly enriched among DEGs in the paternal transcriptome of early two-cell embryos. (**C**) TFs with targets enriched among DEGs in the paternal transcriptome of early two-cell embryos. Colored points indicate significantly enriched TFs after multiple testing correction. (**D**) Schematic illustration showing the intersection between DEGs in the paternal transcriptome of early two-cell embryos and genes that changed in the same direction in sperm from fathers used for IVF to generate hybrid embryos, defining sperm-to-embryo genes. (**E**) Heatmap showing expression of sperm-to-embryo genes in sperm and in the paternal and maternal transcriptomes of early two-cell embryos. (**F**) Stacked bar plot showing the percentage of sperm-to-embryo genes that have the most abundant transcripts in both sperm and the paternal transcriptome of early two-cell embryos. (**G**) TFs whose targets are enriched in sperm-to-embryo genes. Colored points indicate significantly enriched TFs after multiple testing correction. (**H**) Heatmaps showing normalized H3K4me3 ChIP-seq signal for the paternal and maternal genomes in sperm, oocytes, PN5 zygotes, and early and late two-cell embryos. Each row represents a 4-kb region centered on the TSS of sperm-to-embryo or random background genes (±2 kb), ordered by mean ChIP-seq signal. Background genes were randomly chosen from those expressed in early two-cell embryos. Public H3K4me3 ChIP-seq data were derived from Zhang *et al.* (2016)^99^.

We then profiled the transcriptome of sperm from males fed LPD or control diet and identified four DEGs, three downregulated and one upregulated (adjusted P < 0.05, abs(log2FC) > 0.5; Fig. S6B). To relate transcriptomic changes from offspring back to fathers and assess their concordance, we compared the analyses in the paternal transcriptome of early two-cell embryos with log fold changes of corresponding genes in sperm from males used for IVF to generate these embryos (Fig. 3D). These analyses identified 263 “sperm-to-embryo” genes that changed in the same direction in both the paternal transcriptome of embryos and their originated sperm. More than half of sperm-to-embryo genes showed a different direction of expression change in the maternal transcriptome, suggesting parental genome-specific transcriptome effects in the early embryo (Fig. 3E). Analyses of the level of transcripts (including isoforms) showed that for 58% of sperm-to-embryo genes, the most abundant transcript was shared between sperm and early two-cell embryos generated from that sperm (Fig. 3F). This is consistent with the possibility that these transcripts may be directly inherited from sperm.

TF target enrichment analysis of sperm-to-embryo genes identified a set of genome regulators including chromatin remodelers such as KDM5A, KDM5B, ASH2L and CTCF similar to those observed in paternal DEG set of the early two-cell embryo (Fig. 3G). Further, analyses of chromatin features at transcription start sites (TSSs) of sperm-to-embryo genes, using publicly available H3K4me3 ChIP-seq and newly generated sperm ATAC-seq and DNAme datasets, showed strong H3K4me3 enrichment in sperm and moderate enrichment on the paternal genome of pronuclear 5 (PN5) stage zygotes and early two-cell embryos (Fig. 3H). In late two-cell embryos, H3K4me3 signal increased further and was high at regions flanking TSSs but had a characteristic dip at TSSs themselves, consistent with typical flanking nucleosomes marked by H3K4me3 and nucleosome-depleted TSSs ^42^. Sperm ATAC-seq data showed a sharp enrichment of open chromatin accessibility at TSSs (Fig. S6C). In contrast to the paternal genome, the maternal genome showed no detectable H3K4me3 signal at sperm-to-embryo genes in oocytes, minimal signal in PN5 zygotes and a slight enrichment only in late two-cell embryos (Fig. 3H). Since sperm DNAme at CG-rich TSSs can influence H3K4me3 establishment in the zygote ^47^, we next examined DNAme and CG content at the TSS of sperm-to-embryo genes. We observed sharply defined DNAme boundaries, with high methylation flanking TSSs and local depletion at TSSs themselves (Fig. S6D) that coincided with an enrichment of CG nucleotides within ±100 bp (Fig. S6E).

### Paternal LPD alters the maternal transcriptome of early embryos

We next analyzed the maternal transcriptome of the same set of early two-cell embryos and found that 92 genes are downregulated and 78 genes are upregulated in embryos from LPD-fed fathers (Fig. 4A). A direct comparison of paternal and maternal DEGs identified only one overlapping DEG (Fig. S7A). Maternal DEGs were significantly enriched for metabolic and catabolic pathways, including long-chain fatty acid catabolic processes (Fig. S7B). TF target enrichment analyses of maternal DEGs showed enrichment of several regulators including PPARg and components of the TGF-β signaling pathway, SMAD3 and SMAD4 ^48^ (Fig. S7C). We examined if these maternal DEGs were already expressed in oocytes. Because oocyte transcripts frequently have shortened or depleted poly(A) tails ^49^, we used a publicly available rRNA-depleted RNA-seq dataset to ensure full transcriptome coverage independent of polyadenylation status ^50^. Overlap analyses showed that 69% of maternal DEGs were detected in oocytes (Fig. 4B).

**Figure 4.**
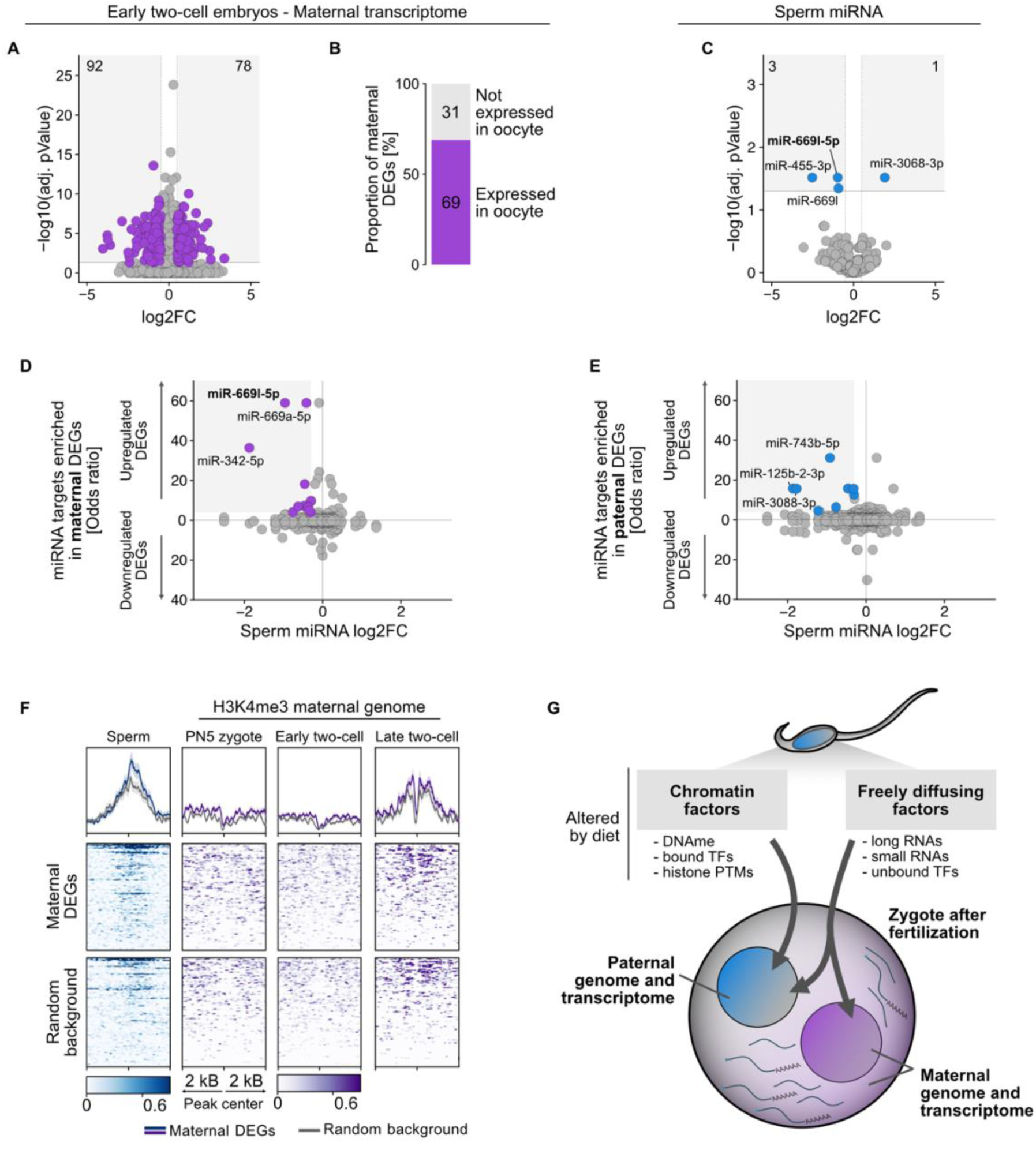
Integrated analyses of maternal transcriptome and sperm miRNAs from LPD-fed C57BL/6J males and chromatin features. (**A**) Volcano plot of DEGs (adjusted *P* < 0.05 and abs(log2FC) > 0.5) in the maternal transcriptome of early two-cell embryos derived from LPD-fed fathers compared to controls. Numbers indicate the count of down- and upregulated DEGs. Control: n = 42, LPD: n = 45. (**B**) Stacked bar plot showing the distribution of maternal DEGs expressed or not expressed in oocytes. Public data are from Wu *et al.* (2020) ^50^. (**C**) Volcano plot of differentially expressed miRNAs (adjusted *P* < 0.05 and abs(log2FC) > 0.5) in sperm from LPD-fed fathers compared to controls. Numbers indicate the count of down- and upregulated miRNAs. Control: n = 9, LPD: n = 9. (**D**) Scatter plot showing sperm miRNA log2FC versus enrichment odds ratio of their predicted targets among maternal DEGs (upregulated, upper half; downregulated, lower half) in early two-cell embryos. Highlighted points indicate miRNAs with reduced expression in sperm whose targets are enriched among upregulated genes in the maternal transcriptome of early two-cell embryos. (**E**) Scatter plot showing sperm miRNA log2FC versus enrichment odds ratio of their predicted targets among paternal DEGs (upregulated, upper half; downregulated, lower half) in early two-cell embryos. Highlighted points indicate miRNAs with reduced expression in sperm whose targets are enriched among upregulated genes in the paternal transcriptome of early two-cell embryos. (**F**) Heatmaps showing normalized H3K4me3 ChIP-seq signal for sperm and the maternal genome in PN5 stage zygotes and early and late two-cell embryos. Each row represents a 4-kb region centered on the TSS of maternal DEGs or random background genes (±2 kb), ordered by mean ChIP-seq signal. Background genes were randomly chosen from those expressed in early two-cell embryos. Public data are from Zhang *et al.* (2016) ^99^. (**G**) Conceptual model illustrating parallel factors that can be transferred from sperm to the paternal and maternal genomes and transcriptomes in the early embryo.

We then analyzed small RNAs in sperm from LPD- and control diet-fed C57BL/6J males and identified four significantly down- or up-regulated microRNAs (miRNA) in response to LPD (Fig. 4C). No change in tRNA or tRNA fragments was detected with the method used (Fig. S7D), though it should be noted that standard small RNA-seq approaches may not capture highly modified tsRNAs and rsRNAs ^51^. Integration of sperm miRNA data with the maternal transcriptome of early two-cell embryos identified a subset of miRNAs with predicted targets enriched among maternal DEGs. miR-669l-5p was significantly downregulated in sperm from LPD-fed males (adjusted P < 0.05) and showed the strongest predicted target enrichment across upregulated maternal DEGs (Fig. 4D). Applying the same integrative analyses to the paternal transcriptome did not detect any differentially expressed sperm miRNAs with enriched targets among paternal DEGs (Fig. 4E).

Analysis of chromatin features at maternal DEGs showed only minimal enrichment of H3K4me3 in sperm and in the maternal genome of early two-cell embryos (Fig. 4F). Likewise, analysis of sperm ATAC-seq, DNAme and H3K27me3 signals did not show any enrichment at these loci (Fig. S7E). Together, these results suggest that paternal LPD is associated with changes in the maternal transcriptome of early two-cell embryos, accompanied by sperm small RNA alterations but no detectable changes in chromatin features at corresponding loci in sperm or the maternal genome of the early embryo.

### Long-term LPD alters gene expression in undifferentiated spermatogonia

Sperm is the product of spermatogenesis initiated by differentiation of spermatogonial stem cells and spermiogenesis in testes, that then matures in the epididymis before being release ^52^. We examined if father-to-offspring effects of paternal LPD involve early stages of spermatogenesis and/or epididymal maturation. We first assessed if the effects can arise in sperm during epididymal transit by feeding C57BL/6J or PWK/PhJ males LPD for 10 days, which is the duration of epididymal sperm maturation ^53^, then mating them with females of the reciprocal strain. 10-day LPD did not affect offspring body weight in either C57BL/6J or PWK/PhJ lineage (Fig. 5, A and B, and Fig. S8, A to F). When LPD treatment was extended to 5 weeks, which spans late stages of spermatogenesis and epididymal transit, there was also no effect on offspring growth, although body weight was slightly reduced in exposed C57BL/6J fathers (−4.5% compared to controls; Fig. 5, A and B, and Fig. S8, G to L). This is in contrast to the effects of 10-week LPD, which fully spans spermatogenesis, spermiogenesis and epididymal transit and induces significant effects in the offspring (Fig. 1, A to G) and fathers (−13.1% weight loss compared to controls; Fig. 1A). These results suggest that early stages of spermatogenesis are involved, pointing to a role of germ cell progenitors.

**Figure 5.**
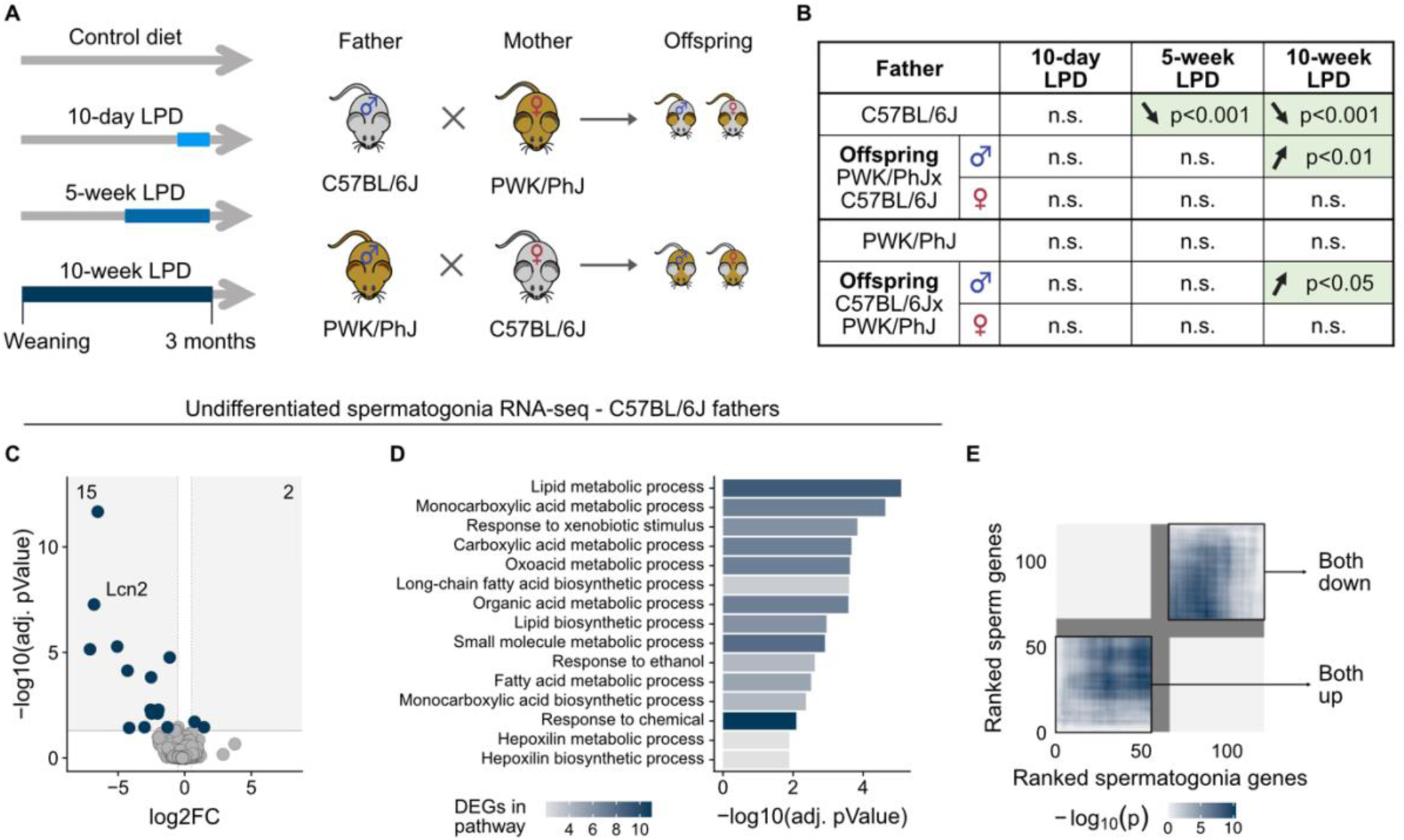
Effects of different LPD duration on body weight and molecular profiling of undifferentiated spermatogonia. (**A**) Schematic illustration showing the timing and duration of LPD exposure in C57BL/6J and PWK/PhJ fathers followed by mating with naïve females of the opposite strain to generate male and female hybrid offspring. (**B**) Summary table of the effects of LPD on body weight growth of C57BL/6J and PWK/PhJ fathers and their male and female hybrid offspring, indicated by *P* values for significant changes. Direction of change is indicated by arrows. (**C**) Volcano plot of DEGs (adjusted *P* < 0.05 and abs(log2FC) > 0.5) in undifferentiated spermatogonia from LPD-exposed C57BL/6J fathers compared to controls. Numbers indicate the count of down- and upregulated DEGs. Control: n = 6, LPD: n = 5. (**D**) Top 15 biological process GO terms most significantly enriched among DEGs in undifferentiated spermatogonia. (**E**) Heatmap showing the RRHO between differential expression analyses of undifferentiated spermatogonia and sperm cells. The color scale indicates the statistical significance of overlap between ranked gene lists in the two datasets. In this RRHO representation, genes in the upper right quadrant correspond to genes downregulated in both cell types (both down), whereas genes in the lower left quadrant correspond to genes upregulated in both cell types (both up).

We thus analyzed spermatogonia by isolating them from adult testes by fluorescence-activated cell sorting (FACS) using MHC I⁻ integrin-α6⁺ Thy1⁺ markers (Fig. S9A) ^54–56^ followed by RNA-seq. Deconvolution analyses using a publicly available single-cell RNA-seq reference of mouse testicular cells showed that samples were primarily attributed to spermatogonial stem cells and spermatogonial progenitor cells, indicating enrichment for undifferentiated spermatogonia (Fig. S9, B to D) ^57^. Differential expression analyses identified 15 downregulated and 2 upregulated genes in response to LPD (adjusted P < 0.05, abs(log2FC) > 0.5; Fig. 5C). These DEGs were enriched in pathways of metabolic processes, particularly lipid metabolism (Fig. 5D). Among the DEGs was the lipid transporter gene *Lcn2*, which notably was reduced in the liver of female offspring from LPD-fed C57BL/6J males ^58^.

To relate spermatogonial cells to sperm, we then compared their respective transcriptome by rank-rank hypergeometric overlap (RRHO) analyses ^59^. Several sets of genes that were downregulated or upregulated in the same direction in both cell stages were identified (Fig. 5E). Analyses of chromatin features of these genes using public histone PTM datasets ^60^ showed that genes downregulated in both spermatogonia and sperm are enriched in H3K27me3 at regions flanking the TSS while upregulated genes are relatively depleted in H3K27me3 (Fig. S10A). This pattern was also observed in sperm, although at lower magnitude (Fig. S10B). Other histone PTMs did not show any consistent enrichment. Together, these results suggest that long-term LPD is associated with transcriptional changes in spermatogonia that overlap with changes in sperm and somatic tissues of exposed males or their offspring.

## Discussion

This study examines the phenotypic consequences and underlying mechanistic basis of paternal dietary restriction in two mouse strains and their offspring. It shows that paternal LPD induces strain-dependent responses in exposed males and sex-specific changes in body weight and gene transcription in their offspring. Using a genetic hybrid design to distinguish paternal and maternal transcriptome in embryos, it shows that transcriptional changes due to paternal LPD are allele-specific in the offspring. These changes are associated with molecular signatures in sperm. Together, the findings suggest that paternal diet can shape early embryonic transcriptome and influence intergenerational outcomes through parallel parent-of-origin molecular routes.

In rodents, LPD is known to modify RNA content and histone PTMs in sperm ^5,13,29^. Further, it is known to alter preimplantation embryos sired by LPD-fed fathers. These embryos show accelerated developmental timing, altered molecular pathways related to metabolism and dysregulation of the endogenous retroelement MERVL (murine endogenous retrovirus-L) ^13,27,46,61^. These findings indicate that paternal diet can influence early stages of embryonic development. Our results significantly extend these findings by focusing on an earlier stage of development before ZGA when embryos are still transcriptionally inactive and rely on parental factors for their development. We show that LPD modifies the transcriptome of preimplantation two-cell embryos, specifically the transcriptome derived from the paternal genome, and that this correlates with changes in the originating sperm. Notably, many genes are altered in the same direction in embryos and sperm. For more than half of them, the same transcript isoforms are similarly represented in both sperm and embryo, consistent with the possibility of direct transcript transfer. Further, these genes are enriched for specific chromatin marks, particularly H3K4me3 on both the sperm genome and the paternal genome of embryos. Consistently, several chromatin remodelers such as *Dot1l* and *Ash1l*, are among the most notable transcriptional regulators that are differentially expressed due to paternal LPD in the embryos. This suggests a link between the transcriptome and chromatin states in sperm and paternal gene regulation in the embryo. Although this link does not prove direct inheritance from sperm to embryo, it may reflect indirect mechanisms, for instance, similar gene regulation across sperm and early embryos or a predisposition of CpG-rich hypomethylated regions in sperm to acquire H3K4me3 during early embryogenesis as described before ^47^. Such regions could act as regulatory sites sensitive to paternal exposure, providing a chromatin-based link between exposure and regulation of the paternal genome in the early embryo. This possibility aligns with previous studies reporting that dietary challenge can alter sperm chromatin and histone PTMs ^19,29,62,63^. Such findings support the idea that environmentally-induced chromatin states in sperm may have regulatory consequences after fertilization. It should be noted that methodological challenges in assessing chromatin accessibility and histone PTMs in mature sperm require caution for the interpretation of chromatin-based associations ^64^.

We observed that paternal LPD is associated with altered expression of maternal transcripts in early embryos, suggesting that paternal exposure influences oocyte-derived RNA populations after fertilization. The differentially expressed maternal genes were enriched for predicted targets of sperm miRNAs that were altered by LPD, pointing to an association between changes in sperm small RNA and maternal transcript expression in the early embryo. In contrast to the paternal transcriptome, maternal DEGs showed no enrichment for H3K4me3 or other chromatin marks assessed in sperm or the early embryo. This suggests that chromatin-associated features are less likely to underlie maternal transcriptional changes. Instead, the observations are consistent with post-transcriptional regulatory processes involving sperm-delivered small RNAs, which respond to paternal exposure and are associated with changes in embryonic gene regulation and offspring phenotypes ^5,13–15,46^. The lack of miRNA target enrichment among paternal DEGs does not exclude the contribution of diffusible sperm-derived factors to paternal gene regulation. This may instead reflect a broader range of regulatory layers through which sperm-derived signals can act on the paternal genome in the embryo. Together, these observations suggest that multiple regulatory features may contribute to the modulation of paternal and maternal transcriptomes in early embryos following paternal LPD.

Our results show that the physiological and molecular consequences of LPD depend on genetic background in fathers and sex in the offspring. Consistent with prior studies reporting strain-dependent responses to nutritional challenges ^65^, we show that C57BL/6J and PWK/PhJ males have distinct sensitivity to LPD, with body weight effects observed only in C57BL/6J males and not PWK/PhJ males. However, PWK/PhJ males had clear transcriptional changes in liver that were nonetheless different from those observed in C57BL/6J males, suggesting distinct molecular responses. The male offspring from both strains had increased body weight while the female offspring had normal weight, consistent with male-specific effects of LPD reported in pure C57BL/6J breeding ^28^. The liver transcriptome was differently altered in male and female offspring, suggesting sex-dependent transcriptional responses to paternal LPD in offspring tissues.

We observed that the effects of paternal LPD on the offspring body weight emerge only after a 10-week exposure while shorter exposure of 10 days or 5 weeks have no detectable impact. This suggests that exposure throughout one full spermatogenic cycle and sperm epididymal transit is necessary to establish intergenerational effects. Small RNA transport during the epididymal maturation can mediate environmental influences on sperm and contribute to offspring phenotypes ^13,66,67^, thus post-testicular processes may play an important role. At the same time, the requirement for prolonged exposure raises the possibility that earlier germ cell populations are affected by LPD. Indeed, undifferentiated spermatogonia had several DEGs that were relevant for the phenotypes e.g. enriched for metabolic processes, and that overlapped with transcriptional changes in sperm. The shared loci between spermatogonia and sperm were associated with H3K27me3 patterns in both cells, a histone PTM linked to chromatin state propagation across cell divisions ^68,69^. Both epididymal maturation and early germ cell stages may contribute to intergenerational effects, potentially in an additive manner rather than via independent pathways. However, the lack of intergenerational effects of short LPD suggests that sustained exposure and the accumulation of molecular changes over time are required. Single-cell RNA-seq of testicular germ cell populations would add to the findings by identifying which cell states are most sensitive to dietary exposure. Such analyses could inform on how long diet-induced transcriptional changes are maintained through successive stages of spermatogenesis.

While this study provides an integrative view of how paternal LPD is associated with early embryonic gene regulation, the analyses are primarily correlative and do not establish causality between specific sperm-derived factors and transcription in the embryo. However, the strength and consistency of the associations identified across multiple regulatory layers, spanning RNA, chromatin accessibility and histone PTMs, substantially narrow down potential molecular mechanisms while reflecting their complexity. The use of two genetically divergent mouse strains is necessary for allele-specific analyses to obtain decisive parental information but introduces phenotyping and molecular complexity. It should also be noted that embryonic transcriptome analyses were performed on IVF-derived embryos while intergenerational phenotypes were obtained from naturally mated cohorts. As sperm are used directly for fertilization without natural mating, IVF minimizes confounding maternal effects and increases the likelihood that transcriptional changes detected in the embryo reflect sperm-associated factors ^70^. However, we cannot exclude that in vitro conditions influence the embryonic transcriptome and may not fully recapitulate the biology underlying the observed offspring phenotypes ^71^. Together, this study highlights how paternal exposure can influence early development through parallel effects on the paternal and maternal genome of the embryo, pointing to a previously uncharacterised way by which sperm may shape offspring development beyond the DNA sequence.

## Materials and methods

### LPD exposure and embryos collection

#### Animal husbandry, diet and body weight analysis

Mice were maintained under a 12 h reverse light/dark cycle in a temperature- and humidity-controlled facility with ad libitum access to food and water. C57BL/6J (C57BL/6JRj, Janvier Labs) and PWK/PhJ (#003715, Jackson Laboratory) breeders were used to generate male pups in-house for LPD exposure. At weaning (21 days of age), males were pseudo-randomly assigned to experimental groups to balance litter origin and body weight across groups. The control diet (Experimental Diet 2000, Kliba Nafag) contained 18.5% crude protein, and the LPD contained 9% crude protein. Males were fed an LPD for 10 weeks (starting at 3 weeks of age), 5 weeks (starting at 8 weeks of age), or 10 days (starting at 11.5 weeks of age). At 3 months of age, LPD exposure was complete and males were either paired with 2-month-old naïve females of the opposite strain maintained on control diet or sacrificed for tissue collection. Males were removed from breeding cages 7-14 days after pairing to exclude post-conception paternal effects. Offspring were maintained on control diet throughout life. Body weight of exposed males was recorded weekly from weaning until breeding or sacrifice at 13 weeks of age, and body weight of offspring was recorded twice weekly from 1 to 3 weeks of age and weekly thereafter until sacrifice at 12 weeks of age. Body weight data were analyzed across time using mixed-effects models to account for nested effects of litter and paternal diet.

#### *In vitro* fertilization and embryo collection

For sperm cryopreservation, caudae epididymides and connected vas deferens were dissected, minced, and placed in a saturated solution of D-(+) raffinose (R7630, Sigma) and skim milk powder (A0830, ITW Reagents), supplemented with 119 mM monothioglycerol (M6145, Sigma). Sperm were allowed to swim out for 10 min, after which tissues were removed and samples were cooled in liquid nitrogen vapor prior to complete freezing in liquid nitrogen and storage at −80 °C. Before IVF, frozen sperm were thawed and capacitated for 1 h in human tubal fluid (HTF) medium (MR-070-D, Merck). PWK/PhJ females (3–5 weeks of age) were hormonally superovulated by intraperitoneal injection of 50 µL HyperOva (KYD-010-06-EX, CosmoBio), followed 47 h later by injection of 5 IU human chorionic gonadotropin (CG10, Sigma). Females were euthanized by cervical dislocation 13 to 14 h after hCG administration, oviducts were collected, and cumulus–oocyte complexes were released into HTF medium. To minimize confounding effects of oocyte quality and batch variability, oocytes collected from each individual female were split equally and fertilized in parallel with sperm from LPD-fed and control diet–fed males. IVF was performed by incubating capacitated motile sperm with cumulus–oocyte complexes in HTF medium supplemented with 1 mM glutathione (G4251, Merck). After 3 h, oocytes were washed four times in HTF medium to remove sperm and cumulus cells and cultured in HTF medium until collection. Individual early two-cell embryos were collected in PBS 20–22 h after fertilization.

### Tissue processing and sequencing library preparation

#### Liver RNA-seq

From each animal, halves of the caudate and left liver lobes were collected and snap-frozen on dry ice. Central regions from both lobes were homogenized in nuclei extraction buffer using gentleMACS C tubes and the gentleMACS Dissociator (program 4C, Miltenyi Biotec). The filtered homogenate was mixed with TRIzol reagent and stored at −80 °C until RNA extraction using the Direct-zol Microprep Kit (Zymo Research) including on-column DNase I treatment. RNA quality and concentration were assessed using the TapeStation RNA ScreenTape assay (Agilent). Ribosomal RNA-depleted libraries were prepared by BGI (Hong Kong, China) using the DNBSEQ LncRNA-Seq protocol and sequenced on a DNBSEQ platform with paired-end 150 bp reads.

#### Liver ATAC-seq

From each animal, halves of the caudate and left liver lobes were collected and snap-frozen on dry ice. Central regions from both lobes were homogenized in nuclei extraction buffer using gentleMACS C tubes and the gentleMACS Dissociator (program 4C, Miltenyi Biotec). The filtered homogenate was washed with ATAC wash buffer (10 mM Tris-HCl pH 7.5, 10 mM NaCl, 3 mM MgCl₂, 0.1% Tween-20) to remove non-nuclear material, and nuclei were counted. A total of 50,000 nuclei were subjected to Tn5 transposition at 37 °C for 30 min following the Omni-ATAC protocol ^72^. Transposed DNA was purified using the MinElute Reaction Cleanup Kit (Qiagen). Libraries were generated using the NEBNext High-Fidelity 2× PCR Master Mix (New England Biolabs) with unique index combinations and amplified for 15 PCR cycles. Library concentration and fragment size distribution were assessed using the TapeStation High Sensitivity D1000 ScreenTape assay (Agilent). Equimolar library pools were sequenced on an Illumina NovaSeq X Plus platform with paired-end 150 bp reads. An initial sequencing run was used to assess read distribution, after which libraries were rebalanced and sequenced to completion. Reads from both runs were merged prior to downstream analysis.

#### Single-embryo RNA-seq

Single-embryo RNA-seq libraries were prepared using the SMART-Seq mRNA LP (with UMIs) kit (Takara Bio). cDNA was amplified for 18 PCR cycles and assessed for concentration using the TapeStation D5000 ScreenTape assay (Agilent). Libraries were prepared from balanced cDNA inputs and amplified for an additional 13 PCR cycles. Library fragment size distribution and concentration were assessed using the TapeStation D5000 ScreenTape assay (Agilent). Equimolar library pools were sequenced on an Illumina NovaSeq X Plus platform with paired-end 150 bp reads. An initial sequencing run was used to assess read distribution, after which libraries were rebalanced and sequenced to completion. Reads from both runs were merged prior to downstream analysis.

#### Sperm rRNA-depleted long RNA-seq

Caudae epididymides and connected vas deferens were dissected, minced, and incubated in M2 medium (M7167, Sigma-Aldrich) to allow sperm to swim out. Motile sperm were enriched by swim-up at 37 °C for 1 h, filtered, washed, and stored in PBS at −80 °C. For RNA extraction, sperm were lysed by combined mechanical and chemical disruption using 0.5 mm stainless steel beads in TRIzol reagent supplemented with Tris(2-carboxyethyl)phosphine hydrochloride (TCEP; 646547, Merck) ^73^, with homogenization performed using a TissueLyser II (Qiagen). RNA was isolated by chloroform extraction, and the aqueous phase was diluted 1:1 with 100% ethanol before purification using the RNA Clean & Concentrator kit (Zymo Research) including on-column DNase I treatment. Ribosomal RNA-depleted libraries were prepared using the SMARTer Stranded Total RNA-Seq Kit v3 – Pico Input Mammalian (Takara Bio) with 4 ng input RNA, 2 min 30 s fragmentation at 94 °C, and 12 PCR2 cycles. Library concentration and fragment size distribution were assessed using the Qubit ds HS assay (ThermoFisher Scientific) and the TapeStation D1000 High Sensitivity ScreenTape assay (Agilent). An equimolar library pool was sequenced on a DNBSEQ platform with paired-end 150 bp reads.

#### Sperm small RNA-seq

Caudae epididymides and connected vas deferens were dissected, minced, and incubated in M2 medium (M7167, Sigma-Aldrich) to allow sperm to swim out. Motile sperm were enriched by swim-up at 37 °C for 1 h, filtered, washed, and stored in PBS at −80 °C. For RNA extraction, sperm were lysed by combined mechanical and chemical disruption using 0.5 mm stainless steel beads in TRIzol reagent supplemented with TCEP (646547, Merck) ^73^, with homogenization performed using a TissueLyser II (Qiagen). RNA was isolated by chloroform extraction, and the aqueous phase was diluted 1:1.5 with 100% ethanol before purification using the RNA Clean & Concentrator kit (Zymo Research) including on-column DNase I treatment. Small RNA-seq libraries were prepared using the NEXTFLEX Small RNA Sequencing Kit v4 (Revvity) with 12.5 ng input RNA. Libraries were amplified for 20 PCR cycles, and library concentration and fragment size distribution were assessed using the Qubit ds HS assay (ThermoFisher Scientific) and the TapeStation D1000 High Sensitivity ScreenTape assay (Agilent). Libraries were pooled at equal molarity and sequenced on an Illumina NextSeq 2000 platform using an XLEAP P2 flow cell with single-end 100 bp reads.

#### Sperm ATAC-seq

Caudae epididymides and connected vas deferens were dissected, minced, and incubated in M2 medium (M7167, Sigma-Aldrich) to allow sperm to swim out. Motile sperm were enriched by swim-up at 37 °C for 1 h, filtered, washed, and counted. A total of 100,000 sperm were lysed on ice in lysis buffer (10 mM Tris-HCl pH 7.5, 10 mM NaCl, 3 mM MgCl₂, 0.1% Tween-20, 0.1% NP-40, 0.01% digitonin) to extract nuclei. Isolated nuclei were subjected to Tn5 transposition at 37 °C for 30 min following the Omni-ATAC protocol ^72^. Transposed DNA was purified using the MinElute Reaction Cleanup Kit (Qiagen). Libraries were generated using the NEBNext High-Fidelity 2× PCR Master Mix (New England Biolabs) with unique index combinations and amplified for 15 PCR cycles. Library concentration and fragment size distribution were assessed using the TapeStation High Sensitivity D1000 ScreenTape assay (Agilent). Equimolar library pools were sequenced on an Illumina NovaSeq 6000 platform with paired-end 50 bp reads.

#### Sperm enzymatic methyl (EM)-seq

Caudae epididymides and connected vas deferens were dissected, minced, and incubated in M2 medium (M7167, Sigma-Aldrich) to allow sperm to swim out. Motile sperm were enriched by swim-up at 37 °C for 1 h, filtered, washed, and stored at −80 °C. For DNA extraction, sperm were lysed by combined mechanical and chemical disruption using 0.5 mm stainless steel beads in TRIzol reagent supplemented with TCEP, with homogenization performed using a TissueLyser II (Qiagen). Lysates were treated with Proteinase K and RNase A, and DNA was extracted using the DNeasy Blood and Tissue Kit (Qiagen) following the manufacturer’s protocol for “Purification of total DNA from animal blood or cells,” with minor modifications. At step 6, 650 µL buffer AW2 was used instead of 500 µL, and the buffer was incubated on the spin column for 5 min before centrifugation. An additional wash with 500 µL buffer AW2 was performed, followed by a final wash with 650 µL of 70% ethanol. DNA concentration was measured using the Qubit ds HS assay (ThermoFisher Scientific), and sample purity was assessed by NanoDrop measurements. For each sample, 50 ng of DNA was used by the Functional Genomics Center Zürich (Zurich, Switzerland) to construct libraries using the NEBNext Enzymatic Methyl-seq Kit (New England Biolabs) according to the manufacturer’s protocol. Libraries were sequenced on an Illumina NovaSeq X Plus platform with paired-end 150 bp reads.

#### Undifferentiated spermatogonia RNA-seq

Both testes were dissected and the tunica albuginea was removed. Seminiferous tubules were digested in 0.5 mg/mL collagenase (C7657, Sigma) prepared in Goni-MEM (phenol-free DMEM supplemented with 1× non-essential amino acids [11140050, Life Technologies], 100 U/mL penicillin–streptomycin [15140122, Gibco], 1 mM sodium pyruvate [11360039, Life Technologies], and 168 µL/L sodium lactate) for 10–12 min at 35 °C. After 5 min sedimentation on ice, collagenase was removed and tubules were incubated in 0.05% trypsin–EDTA (25300054, Gibco) and DNase I (DN25, Sigma) for 5–10 min. Digestion was stopped by addition of fetal calf serum and DNase I, and the cell suspension was filtered through a 40 µm strainer to remove undigested material. Cells were pelleted at 400 × g for 5 min at 4 °C and resuspended in DPBS-S (DPBS supplemented with 1% FCS, 10 mM HEPES, 1 mM pyruvate, 1 mg/mL glucose, and 1× penicillin–streptomycin). The suspension was layered onto a 30% Percoll solution and centrifuged at 600 × g for 8 min at 4 °C without braking. Cells were washed once in DPBS-S and incubated sequentially with primary and secondary antibodies (Table 1), with DPBS-S washes (600 × g, 5 min, 4 °C) between steps. During incubation with the secondary antibody, cells were stained with LIVE/DEAD Fixable Far Red stain (L34973, Invitrogen). After the final wash, cells were pelleted and resuspended in Goni-MEM for sorting. Undifferentiated spermatogonia were sorted using a BD FACSAria III, gating for live MHC I⁻ integrin-α6⁺ Thy1⁺ cells as described previously ^54–56^ (Fig. S9A). Between 5,000 and 8,500 cells per male were collected into Buffer RLT Plus (1053393, Qiagen) supplemented with 1 mM 2-mercaptoethanol (21985023, Life Technologies), snap-frozen on dry ice, and stored at −80 °C.

**Table 1:** List of antibodies used for FACS-based spermatogonia collection.

| Antibody | Company - Cat# | Primary/Secondary | Dilution |
| --- | --- | --- | --- |
| Biotin-conjugated Rat anti-human CD49f (integrin-α6) | Biolegend - 313603 | Primary, conjugated | 1:100 |
| PE Rat anti-mouse CD90.2 (Thy-1.2) | BD Biosciences -553014 | Primary, conjugated | 1:100 |
| BV421 mouse anti-mouse beta1-microglobulin clone S19.8 (MHC I) | BD Biosciences -744802 | Primary | 1:100 |
| Streptavidin Alexa Fluor 488 | Invitrogen - S32354 | Secondary | 1:200 |
| Live/Dead fixable far red | Invitrogen - L34973 |  | 1:1000 |

RNA was extracted using the RNeasy Plus Micro Kit (Qiagen). Ribosomal RNA-depleted libraries were prepared using the SMARTer Stranded Total RNA-Seq Kit v3 – Pico Input Mammalian (Takara Bio) with 10 ng input RNA, 4 min fragmentation at 94 °C, and 10 PCR2 cycles. Library concentration and fragment size distribution were assessed using the Qubit ds HS assay (ThermoFisher Scientific) and the TapeStation D1000 High Sensitivity ScreenTape assay (Agilent). An equimolar library pool was sequenced on an Illumina NovaSeq X Plus platform with paired-end 150 bp reads.

### Bioinformatic analyses

#### Liver RNA-seq data processing and analyses

Raw sequencing reads were trimmed using trimGalore (version 0.6.7) with parameters -q 30, --length 30, and --stringency 2, and quality was assessed using FastQC (version 0.12.1) ^74^ and FastQ Screen (version 0.15.2) ^75^. Allele-specific reads of hybrid offspring were identified by aligning reads to a SNP-masked mm10 reference genome, in which PWK/PhJ SNP positions were N-masked to minimize reference mapping bias, with STAR (version 2.7.10b) ^76^, followed by assignment of allelic origin using SNPsplit (version 0.5.0) ^77^ with --paired and a PWK/PhJ SNP list. Transcript expression was quantified against the mm10 reference genome using Salmon (version 1.10.2) ^78^ with -l A and --validateMappings, and gene-level counts were obtained using TxImport (version 1.22.0) ^79^. Gene counts of offspring samples were merged per father to account for nested experimental design. Differential expression analysis was performed using edgeR (version 3.36.0) ^80^ after removal of unwanted variation using surrogate variable analysis ^81^ implemented via SEtools::svacor (version 1.23.1) with method = “svaseq”. GO biological process enrichment analysis was performed using gprofiler2 (version 0.2.3) ^82^.

#### Sperm and undifferentiated spermatogonia data processing and analyses

Read pairs were tagged with unique molecular identifiers (UMIs) using umi_tools extract (UMItools version 1.1.4) with parameters --bc-pattern NNNNNNNNCCCCCC and --extract-method=string. Raw sequencing reads were trimmed using trimGalore (version 0.6.7) with parameters -q 30, --length 30, and --stringency 2, and quality was assessed using FastQC (version 0.12.1) ^74^ and FastQ Screen (version 0.15.2) ^75^. Residual ribosomal reads were removed using sortMeRNA (version 4.3.6) ^83^ with rRNA reference databases and --fastx option. For UMI-based deduplication, reads were aligned to the mm10 reference genome using STAR (version 2.7.10b) ^76^, deduplicated using umi_tools dedup (UMItools version 1.1.4) with --paired and --buffer-whole-contig, and deduplicated reads were converted back to FASTQ format using samtools fastq (SAMtools version 1.17) ^84^.

Transcript expression was quantified against the mm10 reference genome using Salmon (version 1.10.2)^78^ with -l A and --validateMappings, and gene-level counts were obtained using TxImport (version 1.22.0) ^79^. Differential expression analysis was performed using edgeR (version 3.36.0) ^80^ after removal of unwanted variation using surrogate variable analysis ^81^, implemented via SEtools::svacor (version 1.23.1) with method = “svaseq”. GO biological process enrichment analysis was performed using gprofiler2 (version 0.2.3) ^82^. Overlap analysis between gene expression signatures of undifferentiated spermatogonia and sperm was performed using the rrho2 package (version 1.0) ^59^.

#### Single-embryo RNA-seq data processing

Read pairs were screened for the presence of UMIs by detecting 5′ UMI recognition sequences, appending the extracted UMI sequence to read pair identifiers, and removing UMI recognition sequences from the reads. Raw sequencing reads were trimmed using trimGalore (version 0.6.7) with parameters -q 30, --length 30, and --stringency 2, and quality was assessed using FastQC (version 0.12.1) ^74^ and FastQ Screen (version 0.15.2) ^75^. Allele-specific reads of hybrid offspring were identified by aligning reads to a SNP-masked mm10 reference genome, in which PWK/PhJ SNP positions were N-masked to minimize reference mapping bias, with STAR (version 2.7.10b) ^76^, followed by assignment of allelic origin using SNPsplit (version 0.5.0) ^77^ with --paired and a PWK/PhJ SNP list. For UMI-based deduplication, reads lacking UMIs were removed, remaining reads were aligned to the mm10 reference genome using STAR (version 2.7.10b) ^76^, deduplicated using umi_tools dedup (UMItools version 1.1.4) with --paired and --buffer-whole-contig, and deduplicated reads were converted back to FASTQ format using samtools fastq (SAMtools version 1.17) ^84^. Transcript expression was quantified separately for combined, paternal, or maternal read sets against the mm10 reference genome using Salmon (version 1.10.2) ^78^ with -l A and --validateMappings, and gene-level counts were obtained using TxImport (version 1.22.0) ^79^.

Differential expression analysis was performed using edgeR (version 3.36.0) ^80^ after removal of unwanted variation using surrogate variable analysis ^81^ implemented via SEtools::svacor (version 1.23.1) with method = “svaseq”. The final model additionally included mother (female oocyte donor) as a covariate to account for differences between oocyte donors. GO biological process enrichment analysis was performed using gprofiler2 (version 0.2.3) ^82^. TF and miRNA target enrichment analyses were performed using Enrichr ^85^ by submitting lists of DEGs with background genes corresponding to all genes included in the respective differential expression analysis. TF with enriched targets were selected from the ChEA 2022 database ^86^, and miRNAs with enriched targets were selected from the miRTarBase 2017 database ^87^.

#### Liver ATAC-seq data processing and analyses

Raw sequencing reads were trimmed using trimGalore (version 0.6.7) with parameters -q 30, --length 30, and --stringency 2, and quality was assessed using FastQC (version 0.12.1) ^74^ and FastQ Screen (version 0.15.2) ^75^. Reads were aligned to the mm10 reference genome using Bowtie2 (version 2.5.1) ^88^ with parameters -X 2000, --end-to-end, and --very-sensitive. Duplicate reads, mitochondrial reads, and reads overlapping ENCODE blacklist regions were removed^89^. Reads were shifted to account for Tn5 integration bias using alignmentSieve (deepTools version 3.5.4) ^90^ with --ATACshift option, and nucleosome-free fragments were selected by removing fragments longer than 147 bp. Nucleosome-free fragments were subjected to peak calling using MACS2 callpeak (version 2.2.9.1) ^91^ with parameters --keep-dup auto, --nomodel, and --nolambda.

Differential accessibility analysis was performed using DiffBind (version 3.4.11) ^92^ by generating a common peak set across all samples, quantifying reads within common peak regions, and performing differential analysis using edgeR (version 3.36.0) ^80^ after removal of unwanted variation using surrogate variable analysis ^81^ implemented via SEtools::svacor (version 1.23.1). For offspring samples, BAM files were merged per father, downsampled to the lowest read count, and used for peak calling prior to differential analysis to account for the nested experimental design. To generate heatmaps, samples within each group were normalized, merged, and converted to bigWig files. DARs were subjected to TF motif enrichment analysis using findMotifsGenome.pl (Homer2) ^93^ with GC content normalization.

#### Sperm small RNA-seq data processing and analyses

Sperm small RNA-seq reads were processed as described previously ^46^. Raw sequencing data were trimmed using trimGalore (version 0.6.10) with parameters -q 20, --length 18, and --max_length 40, and quality was assessed using FastQC (version 0.12.1) ^74^. Trimmed reads were aligned to the mm10 reference genome using bowtie (version 1.3.1) ^94^ with parameters -v 0, -k 1, and --best. BED files were generated from BAM alignments using bamtobed (BEDtools version 2.31.1) ^95^. miRNAs were quantified by intersecting BED files with miRNA annotations from miRbase ^96^. For tRNA quantification, trimmed reads were aligned to the GRCm39 reference genome, and counts were obtained by intersecting reads with mm39 tRNA annotations from the GtRNA database ^97^. Differential expression analysis of miRNAs and tRNAs was performed using edgeR (version 3.36.0) ^80^ after removal of unwanted variation using surrogate variable analysis ^81^ implemented via SEtools::svacor (version 1.23.1) with method = “svaseq”.

#### Sperm ATAC-seq data processing

Raw sequencing data reads were trimmed using trimGalore (version 0.6.7) with parameters -q 30, --length 30, and --stringency 2, and quality was assessed using FastQC (version 0.12.1) ^74^ and FastQ Screen (version 0.15.2) ^75^. Reads were aligned to the mm10 reference genome using Bowtie2 (version 2.5.1) ^88^ with parameters -X 2000, --end-to-end, and --very-sensitive. Duplicate reads, mitochondrial reads, and reads overlapping ENCODE blacklist regions were removed ^89^. To generate heatmaps, replicates were normalized, merged, and converted to bigWig files.

#### Sperm DNAme data processing

Raw sequencing reads were trimmed using trimGalore (version 0.6.7) with parameters -q 30, --length 30, and --stringency 2, and quality was assessed using FastQC (version 0.12.1) ^74^ and FastQ Screen (version 0.15.2) ^75^. Trimmed reads were aligned to the mm10 reference genome and cytosine methylation levels were extracted using Bismark (version 0.24.1) ^98^ with default settings. Read-pair information was completed using fixmate (SAMtools version 1.17), and duplicate reads were marked using markdup (SAMtools version 1.17) ^84^. To generate heatmaps, replicates were downsampled, normalized, merged, and converted to bigWig files.

#### Processing of publicly available data

ChIP-seq datasets were downloaded from Zhang *et al*. (2016) ^99^ (sperm and embryo H3K4me3), Zheng *et al.* (2016) ^60^ (sperm and embryo H3K27me3), Cheng *et al.* (2020) ^100^ (spermatogonia H3K4me3 and H3K27me3), and Yue et al. (2014) ^101^ (Liver H3K4me1, H3K4me3 and H3K27ac). Raw sequencing reads were trimmed using trimGalore (version 0.6.7) with parameters -q 30, --length 30, and --stringency 2, and quality was assessed using FastQC (version 0.12.1) ^74^ and FastQ Screen (version 0.15.2) ^75^. Trimmed reads were aligned to the mm10 reference genome using Bowtie2 (version 2.5.1) ^88^ with parameters -X 2000, --end-to-end, and --very-sensitive. Allele-specific reads of hybrid embryo samples were identified by aligning reads to a SNP-masked mm10 reference genome, in which PWK/PhJ SNP positions were N-masked to minimize reference mapping bias, followed by assignment of allelic origin using SNPsplit (version 0.5.0) ^77^ with --paired and a PWK/PhJ SNP list. For heatmap generation, total mapped reads and allele-specific mapped reads were converted to bigWig files.

Publicly available ATAC-seq datasets were downloaded from Lazar-Contes *et al.* (2025) ^56^ (spermatogonia) and processed as described for ChIP-seq datasets, with the additional removal of duplicate reads, mitochondrial reads, and reads overlapping ENCODE blacklist regions after alignment ^89^.

Publicly available RNA-seq datasets were downloaded from Wu *et al.* (2020) ^50^ (oocyte). Raw sequencing reads were trimmed using trimGalore (version 0.6.7) with parameters -q 30, --length 30, and --stringency 2, and quality was assessed using FastQC (version 0.12.1) ^74^ and FastQ Screen (version 0.15.2) ^75^. Transcript expression was quantified against the mm10 reference genome using Salmon (version 1.10.2) ^78^ with -l A and --validateMappings, and gene-level counts were obtained using TxImport (version 1.22.0) ^79^.

Publicly available DNAme datasets were downloaded from Zoch *et al.* (2020) ^102^ (spermatogonia). Raw sequencing reads were trimmed using trimGalore (version 0.6.7) with parameters -q 30, --length 30, and --stringency 2, and quality was assessed using FastQC (version 0.12.1) ^74^ and FastQ Screen (version 0.15.2) ^75^. Trimmed reads were aligned to the mm10 reference genome and cytosine methylation levels were extracted using Bismark (version 0.24.1) ^98^ with default settings. Read-pair information was completed using fixmate (SAMtools version 1.17) ^84^, and duplicate reads were marked using markdup (SAMtools version 1.17). To generate heatmaps, replicates were downsampled, normalized, merged, and converted to bigWig files.

#### Workflow management and reproducibility

All bioinformatic processing steps described above were implemented and executed using Snakemake ^103^ workflows to ensure reproducibility and consistent dependency management. Separate but related workflows were developed for each dataset to accommodate differences in sequencing modality (e.g., RNA-seq, ATAC-seq, small RNA-seq) and library design (e.g., presence or absence of UMIs), while maintaining a consistent organizational structure across analyses.

## Acknowledgements

We thank Theresa Schöpp, Leonardo Zingler Herrero and Maria Dimitriu for assistance with experiments, Francesca Manuella and Chiara Boscardin for assistance with animal experiments and the organization and maintenance of the animal license and Ellen Jaspers for support with writing and administrative coordination. We thank the staff of the Laboratory Animal Services Center (LASC) including Yvonne Zipfel, Sandra Eppenberger and Alberto Corcoba Garcia for animal care and technical support. We thank the Functional Genomics Center Zurich (FGCZ) including Catharine Aquino, Susanne Kreutzer and Hai Bui, for sequencing and technical assistance. We thank members of the Gapp laboratory at ETH Zurich for sharing their IVF protocol and for providing training. We thank the UZH Cytometry Facility for technical support.

ChatGPT (versions GPT-4o and 5.2) was used as writing assistance for the initial draft of the manuscript, which was subsequently revised and improved manually. ChatGPT (versions GPT-4o and 5.2) was also used for code generation in R for data processing and visualization. This work was supported by the University Zurich; ETH Zurich; the Swiss National Science Foundation [grant no 31003A_175742/1]; ETH grants [ETH-10 15–2 and ETH-17 13–2]; the National Centre of Competence in Research (NCCR) RNA&Disease funded by the Swiss National Science Foundation [grant no 182880/Phase 2 and 205601/Phase 3]; the Hochschulmedizin Flagship Project “STRESS”; the European Union Horizon 2020 Research Innovation Program EarlyCause [grant no 848158]; the Horizon Europe programme Staying Healthy 2021 [grant no 101057390 (HappyMums) and 101057529 (FAMILY)] funded by the Swiss State Secretariat for Education, Research and Innovation (SERI); ERA-NET NEURON [SNSF grant no 3200-0-239949 (EMPATHY)]; the FreeNovation grant from Novartis Forschungsstiftung; and the Escher Family Fund. R. Arzate-Mejia received an ETH Postdoctoral Fellowship [grant no 20-1 FEL-28].

## Authors contribution

LCS: Conceptualization, methodology, investigation, formal analysis, visualization, writing – original draft, writing – review and editing

IBG: Investigation

KU: Software, formal analyses

RGAM: Conceptualization, investigation

IMM: Conceptualization, funding acquisition, supervision, writing – review and editing

## References

1. Perez, M.F., and Lehner, B. (2019). Intergenerational and transgenerational epigenetic inheritance in animals. Nat. Cell Biol. 21, 143–151. 10.1038/s41556-018-0242-9.

2. Short, A.K., Yeshurun, S., Powell, R., Perreau, V.M., Fox, A., Kim, J.H., Pang, T.Y., and Hannan, A.J. (2017). Exercise alters mouse sperm small noncoding RNAs and induces a transgenerational modification of male offspring conditioned fear and anxiety. Transl. Psychiatry 7, e1114–e1114. 10.1038/tp.2017.82.

3. Bohacek, J., and Mansuy, I.M. (2015). Molecular insights into transgenerational non-genetic inheritance of acquired behaviours. Nat. Rev. Genet. 16, 641–652. 10.1038/nrg3964.

4. Watkins, A.J., and Sinclair, K.D. (2014). Paternal low protein diet affects adult offspring cardiovascular and metabolic function in mice. Am. J. Physiol. Heart Circ. Physiol. 306, 1444–1452. 10.1152/ajpheart.00981.2013.

5. Chen, Q., Yan, M., Cao, Z., Li, X., Zhang, Y., Shi, J., Feng, G.H., Peng, H., Zhang, X., Zhang, Y., et al. (2016). Sperm tsRNAs contribute to intergenerational inheritance of an acquired metabolic disorder. Science (1979). 351, 397–400. 10.1126/science.aad7977.

6. Franklin, T.B., Russig, H., Weiss, I.C., Gräff, J., Linder, N., Michalon, A., Vizi, S., and Mansuy, I.M. (2010). Epigenetic transmission of the impact of early stress across generations. Biol. Psychiatry 68, 408–415. 10.1016/j.biopsych.2010.05.036.

7. Zheng, X., Li, Z., Wang, G., Wang, H., Zhou, Y., Zhao, X., Cheng, C.Y., Qiao, Y., and Sun, F. (2021). Sperm epigenetic alterations contribute to inter- and transgenerational effects of paternal exposure to long-term psychological stress via evading offspring embryonic reprogramming. Cell Discov. 7, 1–22. 10.1038/s41421-021-00343-5.

8. Anway, M.D., Cupp, A.S., Uzumcu, N., and Skinner, M.K. (2005). Epigenetic transgenerational actions of endocrine disruptors and male fertility. Science (1979). 308, 1466–1469. 10.1126/science.1108190.

9. Brieño-Enríquez, M.A., García-López, J., Cárdenas, D.B., Guibert, S., Cleroux, E., Děd, L., Hourcade, J.D.D., Pěknicová, J., Weber, M., and Del Mazo, J. (2015). Exposure to endocrine disruptor induces transgenerational epigenetic deregulation of microRNAs in primordial germ cells. PLoS One 10, e0124296. 10.1371/journal.pone.0124296.

10. Argaw-Denboba, A., Schmidt, T.S.B., Di Giacomo, M., Ranjan, B., Devendran, S., Mastrorilli, E., Lloyd, C.T., Pugliese, D., Paribeni, V., Dabin, J., et al. (2024). Paternal microbiome perturbations impact offspring fitness. Nature 629, 652–659. 10.1038/s41586-024-07336-w.

11. Masson, B.A., Kiridena, P., Lu, D., Kleeman, E.A., Reisinger, S.N., Qin, W., Davies, W.J., Muralitharan, R.R., Jama, H.A., Antonacci, S., et al. (2025). Depletion of the paternal gut microbiome alters sperm small RNAs and impacts offspring physiology and behavior in mice. Brain Behav. Immun. 123, 290–305. 10.1016/J.BBI.2024.09.020.

12. Fitz-James, M.H., and Cavalli, G. (2022). Molecular mechanisms of transgenerational epigenetic inheritance. Nat. Rev. Genet. 23, 325–341. 10.1038/s41576-021-00438-5.

13. Sharma, U., Conine, C.C., Shea, J.M., Boskovic, A., Derr, A.G., Bing, X.Y., Belleannee, C., Kucukural, A., Serra, R.W., Sun, F., et al. (2016). Biogenesis and function of tRNA fragments during sperm maturation and fertilization in mammals. Science (1979). 351, 391–396. 10.1126/science.aad6780.

14. Gapp, K., Jawaid, A., Sarkies, P., Bohacek, J., Pelczar, P., Prados, J., Farinelli, L., Miska, E., and Mansuy, I.M. (2014). Implication of sperm RNAs in transgenerational inheritance of the effects of early trauma in mice. Nat. Neurosci. 17, 667–669. 10.1038/nn.3695.

15. Tomar, A., Gomez-Velazquez, M., Gerlini, R., Comas-Armangué, G., Makharadze, L., Kolbe, T., Boersma, A., Dahlhoff, M., Burgstaller, J.P., Lassi, M., et al. (2024). Epigenetic inheritance of diet-induced and sperm-borne mitochondrial RNAs. Nature 630, 720– 727. 10.1038/s41586-024-07472-3.

16. Takahashi, Y., Morales Valencia, M., Yu, Y., Ouchi, Y., Takahashi, K., Shokhirev, M.N., Lande, K., Williams, A.E., Fresia, C., Kurita, M., et al. (2023). Transgenerational inheritance of acquired epigenetic signatures at CpG islands in mice. Cell 186, 715–731.e19. 10.1016/J.CELL.2022.12.047.

17. Jönsson, J., Perfilyev, A., Kugelberg, U., Skog, S., Lindström, A., Ruhrmann, S., Ofori, J.K., Bacos, K., Rönn, T., Öst, A., et al. (2025). Impact of excess sugar on the whole genome DNA methylation pattern in human sperm. Epigenomics 17, 89–104. 10.1080/17501911.2024.2439782.

18. Lismer, A., Shao, X., Dumargne, M.C., Lafleur, C., Lambrot, R., Chan, D., Toft, G., Bonde, J.P., Macfarlane, A.J., Bornman, R., et al. (2024). The association between long-term DDT or DDE exposures and an altered sperm epigenome—a cross-sectional study of Greenlandic Inuit and South African VhaVenda Men. Environ. Health Perspect. 132, 017008. 10.1289/EHP12013.

19. Lismer, A., Dumeaux, V., Lafleur, C., Lambrot, R., Brind’Amour, J., Lorincz, M.C., and Kimmins, S. (2021). Histone H3 lysine 4 trimethylation in sperm is transmitted to the embryo and associated with diet-induced phenotypes in the offspring. Dev. Cell 56, 671–686.e6. 10.1016/J.DEVCEL.2021.01.014.

20. Jung, Y.H., Wang, H.L. V., Ruiz, D., Bixler, B.J., Linsenbaum, H., Xiang, J.F., Forestier, S., Shafik, A.M., Jin, P., and Corces, V.G. (2022). Recruitment of CTCF to an Fto enhancer is responsible for transgenerational inheritance of BPA-induced obesity. Proc. Natl. Acad. Sci. U. S. A. 119, e2214988119. 10.1073/pnas.2214988119.

21. Steg, L.C., and Mansuy, I.M. (2025). A father’s legacy: the sperm epigenome, preimplantation development, and paternal environment. Epigenomics 17, 1267– 1280. 10.1080/17501911.2025.2569301.

22. Fan, X., Tang, D., Liao, Y., Li, P., Zhang, Y., Wang, M., Liang, F., Wang, X., Gao, Y., Wen, L., et al. (2020). Single-cell RNA-seq analysis of mouse preimplantation embryos by third-generation sequencing. PLoS Biol. 18, e3001017. 10.1371/JOURNAL.PBIO.3001017.

23. Xia, W., and Xie, W. (2020). Rebooting the epigenomes during mammalian early embryogenesis. Stem Cell Reports 15, 1158–1175. 10.1016/J.STEMCR.2020.09.005.

24. Ma, X., Fan, Y., Xiao, W., Ding, X., Hu, W., and Xia, Y. (2022). Glufosinate-ammonium induced aberrant histone modifications in mouse sperm are concordant with transcriptome in preimplantation embryos. Front. Physiol. 12, 819856. 10.3389/fphys.2021.819856.

25. Nohara, K., Suzuki, T., Okamura, K., Kawai, T., and Nakabayashi, K. (2025). Acquired sperm hypomethylation by gestational arsenic exposure is re-established in both the paternal and maternal genomes of post-epigenetic reprogramming embryos. Epigenetics and Chromatin 18, 1–14. 10.1186/S13072-025-00569-7.

26. Carone, B.R., Fauquier, L., Habib, N., Shea, J.M., Hart, C.E., Li, R., Bock, C., Li, C., Gu, H., Zamore, P.D., et al. (2010). Paternally induced transgenerational environmental reprogramming of metabolic gene expression in mammals. Cell 143, 1084–1096. 10.1016/J.CELL.2010.12.008.

27. Watkins, A.J., Sirovica, S., Stokes, B., Isaacs, M., Addison, O., and Martin, R.A. (2017). Paternal low protein diet programs preimplantation embryo gene expression, fetal growth and skeletal development in mice. Biochimica et Biophysica Acta (BBA) -Molecular Basis of Disease 1863, 1371–1381. 10.1016/J.BBADIS.2017.02.009.

28. Morgan, H.L., Furse, S., Dias, I.H.K., Shabir, K., Castellanos, M., Khan, I., May, S.T., Holmes, N., Carlile, M., Sang, F., et al. (2022). Paternal low protein diet perturbs inter-generational metabolic homeostasis in a tissue-specific manner in mice. Commun. Biol. 5, 1–12. 10.1038/s42003-022-03914-8.

29. Yoshida, K., Maekawa, T., Ly, N.H., Fujita, S. ichiro, Muratani, M., Ando, M., Katou, Y., Araki, H., Miura, F., Shirahige, K., et al. (2020). ATF7-Dependent Epigenetic Changes Are Required for the Intergenerational Effect of a Paternal Low-Protein Diet. Mol. Cell 78, 445–458.e6. 10.1016/J.MOLCEL.2020.02.028.

30. Madhura, R.J., Varsha, A., Chakraborthy, A., B., M.K., Shetty A., V., and Badanthadka, M. (2023). Protein malnutrition in BALB/C mice: A model mimicking clinical scenario of marasmic-kwashiorkor malnutrition. J. Pharmacol. Toxicol. Methods 119, 107231. 10.1016/J.VASCN.2022.107231.

31. Smith, K., Dennis, K.M.J.H., and Hodson, L. (2025). The ins and outs of liver fat metabolism: The effect of phenotype and diet on risk of intrahepatic triglyceride accumulation. Exp. Physiol. 110, 936–948. 10.1113/EP092001.

32. Xiao, X., Kennelly, J.P., Ferrari, A., Clifford, B.L., Whang, E., Gao, Y., Qian, K., Sandhu, J., Jarrett, K.E., Brearley-Sholto, M.C., et al. (2023). Hepatic nonvesicular cholesterol transport is critical for systemic lipid homeostasis. Nat. Metab. 5, 165–181. 10.1038/s42255-022-00722-6.

33. Pérez-Martí, A., Garcia-Guasch, M., Tresserra-Rimbau, A., Carrilho-Do-Rosário, A., Estruch, R., Salas-Salvadó, J., Martínez-González, M.Á., Lamuela-Raventós, R., Marrero, P.F., Haro, D., et al. (2017). A low-protein diet induces body weight loss and browning of subcutaneous white adipose tissue through enhanced expression of hepatic fibroblast growth factor 21 (FGF21). Mol. Nutr. Food Res. 61, 1600725. 10.1002/MNFR.201600725.

34. Yamada, R., Odamaki, S., Araki, M., Watanabe, T., Matsuo, K., Uchida, K., Kato, T., Ozaki-Masuzawa, Y., and Takenaka, A. (2019). Dietary protein restriction increases hepatic leptin receptor mRNA and plasma soluble leptin receptor in male rodents. PLoS One 14, e0219603. 10.1371/JOURNAL.PONE.0219603.

35. Klemm, S.L., Shipony, Z., and Greenleaf, W.J. (2019). Chromatin accessibility and the regulatory epigenome. Nat. Rev. Genet. 20, 207–220. 10.1038/S41576-018-0089-8.

36. Millán-Zambrano, G., Burton, A., Bannister, A.J., and Schneider, R. (2022). Histone post-translational modifications — cause and consequence of genome function. Nat. Rev. Genet. 23, 563–580. 10.1038/s41576-022-00468-7.

37. Lau, H.H., Ng, N.H.J., Loo, L.S.W., Jasmen, J.B., and Teo, A.K.K. (2018). The molecular functions of hepatocyte nuclear factors – In and beyond the liver. J. Hepatol. 68, 1033– 1048. 10.1016/J.JHEP.2017.11.026.

38. Montaigne, D., Butruille, L., and Staels, B. (2021). PPAR control of metabolism and cardiovascular functions. Nat. Rev. Cardiol. 18, 809–823. 10.1038/s41569-021-00569-6.

39. Huang, Y., He, S., Li, J.Z., Seo, Y.K., Osborne, T.F., Cohen, J.C., and Hobbs, H.H. (2010). A feed-forward loop amplifies nutritional regulation of PNPLA3. Proc. Natl. Acad. Sci. U. S. A. 107, 7892–7897. 10.1073/pnas.1003585107.

40. Xia, M.F., Lin, H.D., Chen, L.Y., Wu, L., Ma, H., Li, Q., Aleteng, Q., Hu, Y., He, W.Y., Gao, J., et al. (2019). The PNPLA3 rs738409 C>G variant interacts with changes in body weight over time to aggravate liver steatosis, but reduces the risk of incident type 2 diabetes. Diabetologia 62, 644–654. 10.1007/S00125-018-4805-X.

41. Hoek-van den Hil, E.F., van Schothorst, E.M., van der Stelt, I., Swarts, H.J.M., van Vliet, M., Amolo, T., Vervoort, J.J.M., Venema, D., Hollman, P.C.H., Rietjens, I.M.C.M., et al. (2015). Direct comparison of metabolic health effects of the flavonoids quercetin, hesperetin, epicatechin, apigenin and anthocyanins in high-fat-diet-fed mice. Genes Nutr. 10, 23. 10.1007/S12263-015-0469-Z.

42. De Dieuleveult, M., Yen, K., Hmitou, I., Depaux, A., Boussouar, F., Dargham, D.B., Jounier, S., Humbertclaude, H., Ribierre, F., Baulard, C., et al. (2016). Genome-wide nucleosome specificity and function of chromatin remodellers in ES cells. Nature 530, 113–116. 10.1038/nature16505.

43. Deng, Q., Ramsköld, D., Reinius, B., and Sandberg, R. (2014). Single-cell RNA-seq reveals dynamic, random monoallelic gene expression in mammalian cells. Science (1979). 343, 193–196. 10.1126/science.1245316.

44. Blanco, M., El Khattabi, L., Gobé, C., Crespo, M., Coulée, M., de la Iglesia, A., Ialy-Radio, C., Lapoujade, C., Givelet, M., Delessard, M., et al. (2023). DOT1L regulates chromatin reorganization and gene expression during sperm differentiation. EMBO Rep. 24. 10.15252/EMBR.202256316.

45. Vann, K.R., Sharma, R., Hsu, C.C., Devoucoux, M., Tencer, A.H., Zeng, L., Lin, K., Zhu, L., Li, Q., Lachance, C., et al. (2025). Structure-function relationship of ASH1L and histone H3K36 and H3K4 methylation. Nat. Commun. 16, 1–15. 10.1038/s41467-025-57556-5.

46. Dura, M., Ranjan, B., Serrano, J.B., Paribeni, R., Paribeni, V., Villacorta, L., Benes, V., Boruc, O., Boskovic, A., and Hackett, J.A. (2025). Embryonic signatures of intergenerational epigenetic inheritance across paternal environments and genetic backgrounds. EMBO Journal 44, 6750–6781. 10.1038/S44318-025-00556-4.

47. Fanourgakis, G., Gaspa-Toneu, L., Komarov, P.A., Papasaikas, P., Ozonov, E.A., Smallwood, S.A., and Peters, A.H.F.M. (2025). DNA methylation modulates nucleosome retention in sperm and H3K4 methylation deposition in early mouse embryos. Nat. Commun. 16, 1–22. 10.1038/s41467-024-55441-1.

48. Zhang, L., Duan, C.J., Binkley, C., Li, G., Uhler, M.D., Logsdon, C.D., and Simeone, D.M. (2004). A Transforming growth factor β-induced Smad3/Smad4 complex directly activates protein kinase A. Mol. Cell. Biol. 24, 2169. 10.1128/MCB.24.5.2169-2180.2004.

49. Lee, K., Cho, K., Morey, R., and Cook-Andersen, H. (2024). An extended wave of global mRNA deadenylation sets up a switch in translation regulation across the mammalian oocyte-to-embryo transition. Cell Rep. 43, 113710. 10.1016/J.CELREP.2024.113710.

50. Wu, D., and Dean, J. (2020). EXOSC10 sculpts the transcriptome during the growth-to-maturation transition in mouse oocytes. Nucleic Acids Res. 48, 5349–5365. 10.1093/NAR/GKAA249.

51. Shi, J., Zhang, Y., Tan, D., Zhang, X., Yan, M., Zhang, Y., Franklin, R., Shahbazi, M., Mackinlay, K., Liu, S., et al. (2021). PANDORA-seq expands the repertoire of regulatory small RNAs by overcoming RNA modifications. Nat. Cell Biol. 23, 424–436. 10.1038/s41556-021-00652-7.

52. Sultana, T., and Sutovsky, P. (2026). A concise overview of mammalian spermatogenesis. Syst. Biol. Reprod. Med. 72, 3–22. 10.1080/19396368.2025.2593339.

53. Clermont, Y. (1972). Kinetics of spermatogenesis in mammals: seminiferous epithelium cycle and spermatogonial renewal. Physiol. Rev. 52, 198–236. 10.1152/PHYSREV.1972.52.1.198.

54. Kubota, H., Avarbock, M.R., and Brinster, R.L. (2004). Culture conditions and single growth factors affect fate determination of mouse spermatogonial stem cells. Biol. Reprod. 71, 722–731. 10.1095/BIOLREPROD.104.029207.

55. Kubota, H., Avarbock, M.R., and Brinster, R.L. (2003). Spermatogonial stem cells share some, but not all, phenotypic and functional characteristics with other stem cells. Proc. Natl. Acad. Sci. U. S. A. 100, 6487–6492. 10.1073/PNAS.0631767100.

56. Lazar-Contes, I., Arzate-Mejia, R.G., Tanwar, D.K., Steg, L.C., Uzel, K., Feudjio, O.U., Crespo, M., Germain, P.-L., and Mansuy, I.M. (2025). Dynamics of transcriptional programs and chromatin accessibility in mouse spermatogonial cells from early postnatal to adult life. Elife 12. 10.7554/ELIFE.91528.

57. Hermann, B.P., Cheng, K., Singh, A., Roa-De La Cruz, L., Mutoji, K.N., Chen, I.C., Gildersleeve, H., Lehle, J.D., Mayo, M., Westernströer, B., et al. (2018). The mammalian spermatogenesis single-cell transcriptome, from spermatogonial stem cells to spermatids. Cell Rep. 25, 1650–1667.e8. 10.1016/J.CELREP.2018.10.026.

58. Jaberi, S. Al, Cohen, A., D’Souza, C., Abdulrazzaq, Y.M., Ojha, S., Bastaki, S., and Adeghate, E.A. (2021). Lipocalin-2: Structure, function, distribution and role in metabolic disorders. Biomedicine & Pharmacotherapy 142, 112002. 10.1016/J.BIOPHA.2021.112002.

59. Plaisier, S.B., Taschereau, R., Wong, J.A., and Graeber, T.G. (2010). Rank–rank hypergeometric overlap: identification of statistically significant overlap between gene-expression signatures. Nucleic Acids Res. 38, e169–e169. 10.1093/NAR/GKQ636.

60. Zheng, H., Huang, B., Zhang, B., Xiang, Y., Du, Z., Xu, Q., Li, Y., Wang, Q., Ma, J., Peng, X., et al. (2016). Resetting epigenetic memory by reprogramming of histone modifications in mammals. Mol. Cell 63, 1066–1079. 10.1016/J.MOLCEL.2016.08.032.

61. Morgan, H.L., Eid, N., Holmes, N., Henson, S., Wright, V., Coveney, C., Winder, C., O’Neil, D.M., Dunn, W.B., Boocock, D.J., et al. (2024). Paternal undernutrition and overnutrition modify semen composition and preimplantation embryo developmental kinetics in mice. BMC Biol. 22, 207-. 10.1186/S12915-024-01992-0.

62. Tahiri, I., Llana, S.R., Fos-Domènech, J., Milà-Guash, M., Toledo, M., Haddad-Tóvolli, R., Claret, M., and Obri, A. (2024). Paternal obesity induces changes in sperm chromatin accessibility and has a mild effect on offspring metabolic health. Heliyon 10, e34043. 10.1016/J.HELIYON.2024.E34043.

63. Pepin, A.-S., Jazwiec, P.A., Dumeaux, V., Sloboda, D.M., and Kimmins, S. (2024). Determining the effects of paternal obesity on sperm chromatin at histone H3 lysine 4 tri-methylation in relation to the placental transcriptome and cellular composition. Elife 13. 10.7554/ELIFE.83288.

64. Yin, Q., Yang, C.H., Strelkova, O.S., Wu, J., Sun, Y., Gopalan, S., Yang, L., Dekker, J., Fazzio, T.G., Li, X.Z., et al. (2023). Revisiting chromatin packaging in mouse sperm. Genome Res. 33, 2079–2093. 10.1101/GR.277845.123.

65. Bachmann, A.M., Morel, J.D., El Alam, G., Rodríguez-López, S., Imamura de lima, T., Goeminne, L.J.E., Benegiamo, G., Loric, S., Conti, M., Sleiman, M.B., et al. (2022). Genetic background and sex control the outcome of high-fat diet feeding in mice. iScience 25, 104468. 10.1016/J.ISCI.2022.104468.

66. Conine, C.C., Sun, F., Song, L., Rivera-Pérez, J.A., and Rando, O.J. (2018). Small RNAs gained during epididymal transit of sperm are essential for embryonic development in mice. Dev. Cell 46, 470–480.e3. 10.1016/J.DEVCEL.2018.06.024.

67. Trigg, N.A., and Conine, C.C. (2024). Epididymal acquired sperm microRNAs modify post-fertilization embryonic gene expression. Cell Rep. 43, 114698. 10.1016/J.CELREP.2024.114698.

68. Mikulski, P., Tehrani, S.S.H., Kogan, A., Abdul-Zani, I., Shell, E., James, L., Ryan, B.J., and Jansen, L.E.T. (2025). Heritable maintenance of chromatin modifications confers transcriptional memory of interferon-γ signaling. Nat. Struct. Mol. Biol. 32, 1255–1267. 10.1038/s41594-025-01522-8.

69. Hansen, K.H., Bracken, A.P., Pasini, D., Dietrich, N., Gehani, S.S., Monrad, A., Rappsilber, J., Lerdrup, M., and Helin, K. (2008). A model for transmission of the H3K27me3 epigenetic mark. Nat. Cell Biol. 10, 1291–1300. 10.1038/ncb1787.

70. Bohacek, J., and Mansuy, I.M. (2017). A guide to designing germline-dependent epigenetic inheritance experiments in mammals. Nat. Methods 14, 243–249. 10.1038/nmeth.4181.

71. van der Weijden, V.A., Schmidhauser, M., Kurome, M., Knubben, J., Flöter, V.L., Wolf, E., and Ulbrich, S.E. (2021). Transcriptome dynamics in early in vivo developing and in vitro produced porcine embryos. BMC Genomics 22, 139. 10.1186/S12864-021-07430-7.

72. Corces, M.R., Trevino, A.E., Hamilton, E.G., Greenside, P.G., Sinnott-Armstrong, N.A., Vesuna, S., Satpathy, A.T., Rubin, A.J., Montine, K.S., Wu, B., et al. (2017). An improved ATAC-seq protocol reduces background and enables interrogation of frozen tissues. Nat. Methods 14, 959–962. 10.1038/nmeth.4396.

73. Roszkowski, M., and Mansuy, I.M. (2021). High efficiency RNA extraction fom sperm cells using Guanidinium Thiocyanate supplemented with Tris(2-Carboxyethyl)Phosphine. Front. Cell Dev. Biol. 9, 648274. 10.3389/FCELL.2021.648274.

74. Andrews, S. (2010). FastQC: a quality control tool for high throughput sequence data. https://www.bioinformatics.babraham.ac.uk/projects/fastqc/.

75. Wingett, S.W., and Andrews, S. (2018). FastQ Screen: a tool for multi-genome mapping and quality control. F1000Res. 7, 1338. 10.12688/F1000RESEARCH.15931.2.

76. Dobin, A., Davis, C.A., Schlesinger, F., Drenkow, J., Zaleski, C., Jha, S., Batut, P., Chaisson, M., and Gingeras, T.R. (2013). STAR: ultrafast universal RNA-seq aligner. Bioinformatics 29, 15–21. 10.1093/BIOINFORMATICS/BTS635.

77. Krueger, F., and Andrews, S.R. (2016). SNPsplit: allele-specific splitting of alignments between genomes with known SNP genotypes. F1000Res. 5, 1479. 10.12688/f1000research.9037.2.

78. Patro, R., Duggal, G., Love, M.I., Irizarry, R.A., and Kingsford, C. (2017). Salmon provides fast and bias-aware quantification of transcript expression. Nat. Methods 14, 417–419. 10.1038/nmeth.4197.

79. Soneson, C., Love, M.I., and Robinson, M.D. (2016). Differential analyses for RNA-seq: transcript-level estimates improve gene-level inferences. F1000Res. 4, 1521. 10.12688/f1000research.7563.2.

80. Robinson, M.D., McCarthy, D.J., and Smyth, G.K. (2010). edgeR: a Bioconductor package for differential expression analysis of digital gene expression data. Bioinformatics 26, 139–140. 10.1093/BIOINFORMATICS/BTP616.

81. Leek, J.T., Johnson, W.E., Parker, H.S., Jaffe, A.E., and Storey, J.D. (2012). The sva package for removing batch effects and other unwanted variation in high-throughput experiments. Bioinformatics 28, 882–883. 10.1093/BIOINFORMATICS/BTS034.

82. Kolberg, L., Raudvere, U., Kuzmin, I., Adler, P., Vilo, J., and Peterson, H. (2023). g:Profiler—interoperable web service for functional enrichment analysis and gene identifier mapping (2023 update). Nucleic Acids Res. 51, W207–W212. 10.1093/NAR/GKAD347.

83. Kopylova, E., Noé, L., and Touzet, H. (2012). SortMeRNA: fast and accurate filtering of ribosomal RNAs in metatranscriptomic data. Bioinformatics 28, 3211–3217. 10.1093/BIOINFORMATICS/BTS611.

84. Danecek, P., Bonfield, J.K., Liddle, J., Marshall, J., Ohan, V., Pollard, M.O., Whitwham, A., Keane, T., McCarthy, S.A., and Davies, R.M. (2021). Twelve years of SAMtools and BCFtools. Gigascience 10, 1–4. 10.1093/GIGASCIENCE/GIAB008.

85. Kuleshov, M. V., Jones, M.R., Rouillard, A.D., Fernandez, N.F., Duan, Q., Wang, Z., Koplev, S., Jenkins, S.L., Jagodnik, K.M., Lachmann, A., et al. (2016). Enrichr: a comprehensive gene set enrichment analysis web server 2016 update. Nucleic Acids Res. 44, W90–W97. 10.1093/NAR/GKW377.

86. Keenan, A.B., Torre, D., Lachmann, A., Leong, A.K., Wojciechowicz, M.L., Utti, V., Jagodnik, K.M., Kropiwnicki, E., Wang, Z., and Ma’ayan, A. (2019). ChEA3: transcription factor enrichment analysis by orthogonal omics integration. Nucleic Acids Res. 47, W212–W224. 10.1093/NAR/GKZ446.

87. Chou, C.H., Chang, N.W., Shrestha, S., Hsu, S. Da, Lin, Y.L., Lee, W.H., Yang, C.D., Hong, H.C., Wei, T.Y., Tu, S.J., et al. (2016). miRTarBase 2016: updates to the experimentally validated miRNA-target interactions database. Nucleic Acids Res. 44, D239–D247. 10.1093/NAR/GKV1258.

88. Langmead, B., and Salzberg, S.L. (2012). Fast gapped-read alignment with Bowtie 2. Nature 9, 357–359. 10.1038/nmeth.1923.

89. Amemiya, H.M., Kundaje, A., and Boyle, A.P. (2019). The ENCODE blacklist: identification of problematic regions of the genome. Sci. Rep. 9, 1–5. 10.1038/s41598-019-45839-z.

90. Ramírez, F., Ryan, D.P., Grüning, B., Bhardwaj, V., Kilpert, F., Richter, A.S., Heyne, S., Dündar, F., and Manke, T. (2016). deepTools2: a next generation web server for deep-sequencing data analysis. Nucleic Acids Res. 44, W160–W165. 10.1093/NAR/GKW257.

91. Zhang, Y., Liu, T., Meyer, C.A., Eeckhoute, J., Johnson, D.S., Bernstein, B.E., Nussbaum, C., Myers, R.M., Brown, M., Li, W., et al. (2008). Model-based analysis of ChIP-Seq (MACS). Genome Biol. 9, 1–9. 10.1186/GB-2008-9-9-R137.

92. Ross-Innes, C.S., Stark, R., Teschendorff, A.E., Holmes, K.A., Ali, H.R., Dunning, M.J., Brown, G.D., Gojis, O., Ellis, I.O., Green, A.R., et al. (2012). Differential oestrogen receptor binding is associated with clinical outcome in breast cancer. Nature 481, 389–393. 10.1038/nature10730.

93. Heinz, S., Benner, C., Spann, N., Bertolino, E., Lin, Y.C., Laslo, P., Cheng, J.X., Murre, C., Singh, H., and Glass, C.K. (2010). Simple combinations of lineage-determining transcription factors prime cis-regulatory elements required for macrophage and B cell identities. Mol. Cell 38, 576–589. 10.1016/J.MOLCEL.2010.05.004.

94. Langmead, B., Trapnell, C., Pop, M., and Salzberg, S.L. (2009). Ultrafast and memory-efficient alignment of short DNA sequences to the human genome. Genome Biol. 10, 1–10. 10.1186/GB-2009-10-3-R25.

95. Quinlan, A.R., and Hall, I.M. (2010). BEDTools: a flexible suite of utilities for comparing genomic features. Bioinformatics 26, 841–842. 10.1093/BIOINFORMATICS/BTQ033.

96. Kozomara, A., Birgaoanu, M., and Griffiths-Jones, S. (2019). miRBase: from microRNA sequences to function. Nucleic Acids Res. 47, D155–D162. 10.1093/NAR/GKY1141.

97. Chan, P.P., and Lowe, T.M. (2016). GtRNAdb 2.0: an expanded database of transfer RNA genes identified in complete and draft genomes. Nucleic Acids Res. 44, D184–D189. 10.1093/NAR/GKV1309.

98. Krueger, F., and Andrews, S.R. (2011). Bismark: a flexible aligner and methylation caller for Bisulfite-Seq applications. Bioinformatics 27, 1571–1572. 10.1093/BIOINFORMATICS/BTR167.

99. Zhang, B., Zheng, H., Huang, B., Li, W., Xiang, Y., Peng, X., Ming, J., Wu, X., Zhang, Y., Xu, Q., et al. (2016). Allelic reprogramming of the histone modification H3K4me3 in early mammalian development. Nature 537, 553–557. 10.1038/nature19361.

100. Cheng, K., Chen, I.C., Cheng, C.H.E., Mutoji, K., Hale, B.J., Hermann, B.P., Geyer, C.B., Oatley, J.M., and McCarrey, J.R. (2020). Unique epigenetic programming distinguishes regenerative spermatogonial stem cells in the developing mouse testis. iScience 23, 101596. 10.1016/J.ISCI.2020.101596/ATTACHMENT/C01C8006-C0C4-4E5F-95D2-2C4ACB4630DF/MMC5.XLSX.

101. Yue, F., Cheng, Y., Breschi, A., Vierstra, J., Wu, W., Ryba, T., Sandstrom, R., Ma, Z., Davis, C., Pope, B.D., et al. (2014). A comparative encyclopedia of DNA elements in the mouse genome. Nature 515, 355–364. 10.1038/nature13992.

102. Zoch, A., Auchynnikava, T., Berrens, R. V., Kabayama, Y., Schöpp, T., Heep, M., Vasiliauskaitė, L., Pérez-Rico, Y.A., Cook, A.G., Shkumatava, A., et al. (2020). SPOCD1 is an essential executor of piRNA-directed de novo DNA methylation. Nature 584, 635– 639. 10.1038/s41586-020-2557-5.

103. Mölder, F., Jablonski, K.P., Letcher, B., Hall, M.B., Tomkins-Tinch, C.H., Sochat, V., Forster, J., Lee, S., Twardziok, S.O., Kanitz, A., et al. (2021). Sustainable data analysis with Snakemake. F1000Res. 10, 33. 10.12688/F1000RESEARCH.29032.2.

