## Supplementary figures for "Paternal diet shapes paternal and maternal transcriptomes in the early embryo through parallel sperm-associated mechanisms"

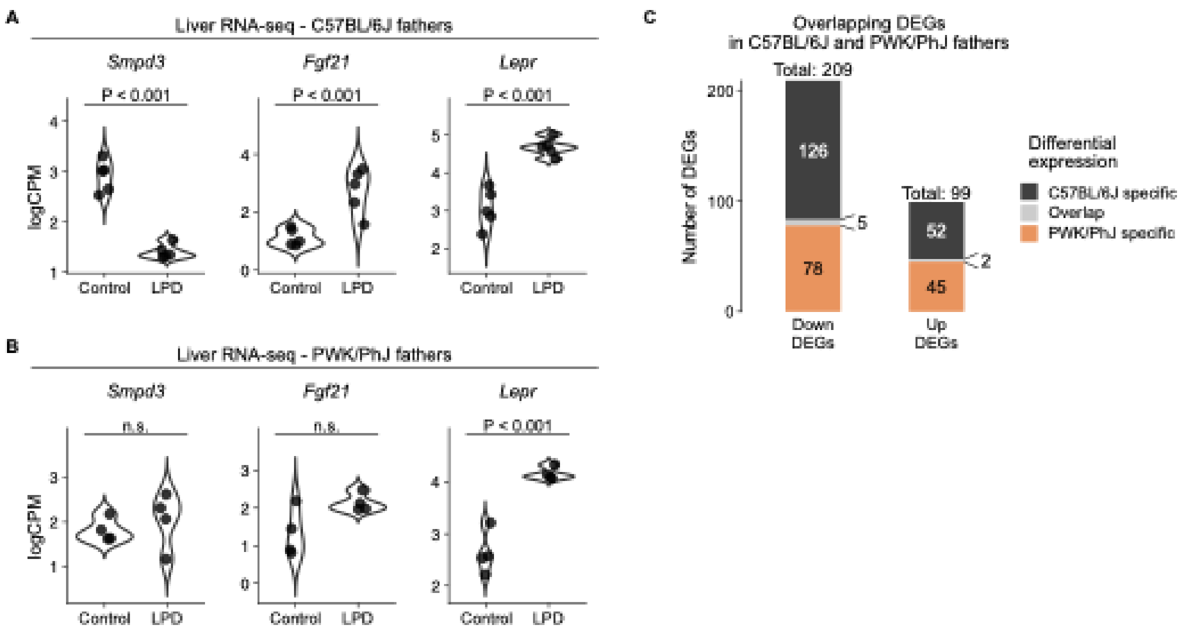


**Figure S1. Analyses of overlapping gene expression in C57BL/6J and PWK/PhJ males fed LPD.** (**A** and **B**) Violin plots showing expression of selected genes in control and LPD groups for (**A**) C57BL/6J and (**B**) PWK/PhJ males. *P* values are derived from differential expression analyses. (**C**) Stacked bar plots showing the overlap of down- and upregulated DEGs between C57BL/6J and PWK/PhJ males.

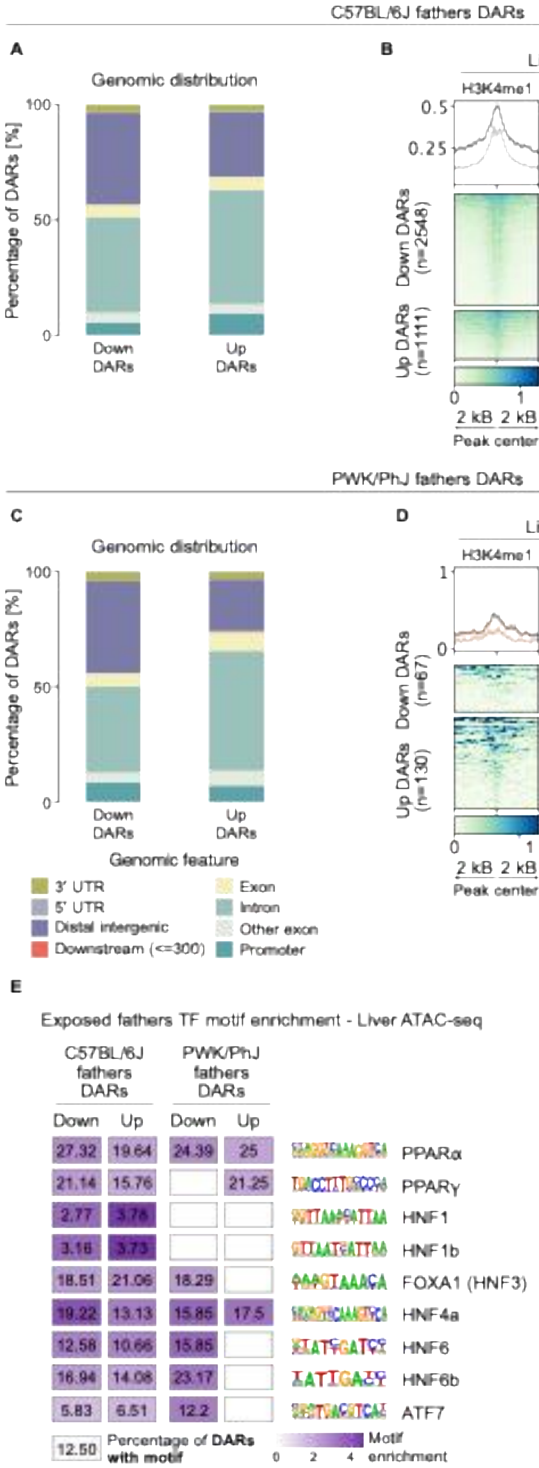

C57BL/6J fathers DARs
A
Genomic distribution
Percentage of DARs [%]
100
50
0
Down DARs
Up DARs
B
Liver chromatin
H3K4me1
H3K4me3
H3K27ac
Down
Up
0.5
0.25
Down DARs (n=2548)
Up DARs (n=1111)
0
1
2 kB
2 kB
Peak center
PWK/PhJ fathers DARs
C
Genomic distribution
Percentage of DARs [%]
100
50
0
Down DARs
Up DARs
D
Liver chromatin
H3K4me1
H3K4me3
H3K27ac
Down
Up
1
0
Down DARs (n=67)
Up DARs (n=130)
2 kB
2 kB
Peak center
Genomic feature
3' UTR
5' UTR
Distal intergenic
Downstream (<=300)
Exon
Intron
Other exon
Promoter
E
Exposed fathers TF motif enrichment - Liver ATAC-seq
C57BL/6J fathers DARs
PWK/PhJ fathers DARs
Down
Up
Down
Up
27.32
19.64
24.39
25
PPARα
21.14
15.76
21.25
PPARγ
2.77
3.76
HNF1
3.16
3.73
HNF1b
18.51
21.06
18.29
FOXA1 (HNF3)
19.22
13.13
15.85
17.5
HNF4a
12.58
10.66
15.85
HNF6
16.94
14.08
23.17
HNF6b
5.83
6.51
12.2
ATF7
12.50
Percentage of DARs with motif
0
2
4
Motif enrichment

**Figure S2. Analyses of DARs in C57BL/6J and PWK/PhJ males fed LPD**. (**A** and **B**) Genomic annotation of DARs identified in C57BL/6J males. (**A**) Stacked bar plot showing the genomic distribution of DARs. (**B**) Heatmaps showing normalized H3K4me1, H3K4me3, and H3K27ac ChIP-seq signal in liver. Each row represents a 4-kb region centered on the DAR midpoint (±2 kb). Public data are from Yue *et al.* (2014) [101]. (**C** and **D**) Genomic annotation of DARs identified in PWK/PhJ males. (**C**) Stacked bar plot showing the genomic distribution of DARs. (**D**) Heatmaps showing normalized H3K4me1, H3K4me3, and H3K27ac ChIP-seq signal in liver. Each row represents a 4-kb region centered on the DAR midpoint (±2 kb). (**E**) Enrichment of metabolic TF motifs in DARs identified in C57BL/6J and PWK/PhJ males fed LPD. Only motifs with significant enrichment after multiple testing correction are shown. The color scale indicates fold enrichment, and numbers indicate the percentage of DARs containing each motif. Empty panels indicate no significant TF motif enrichment.

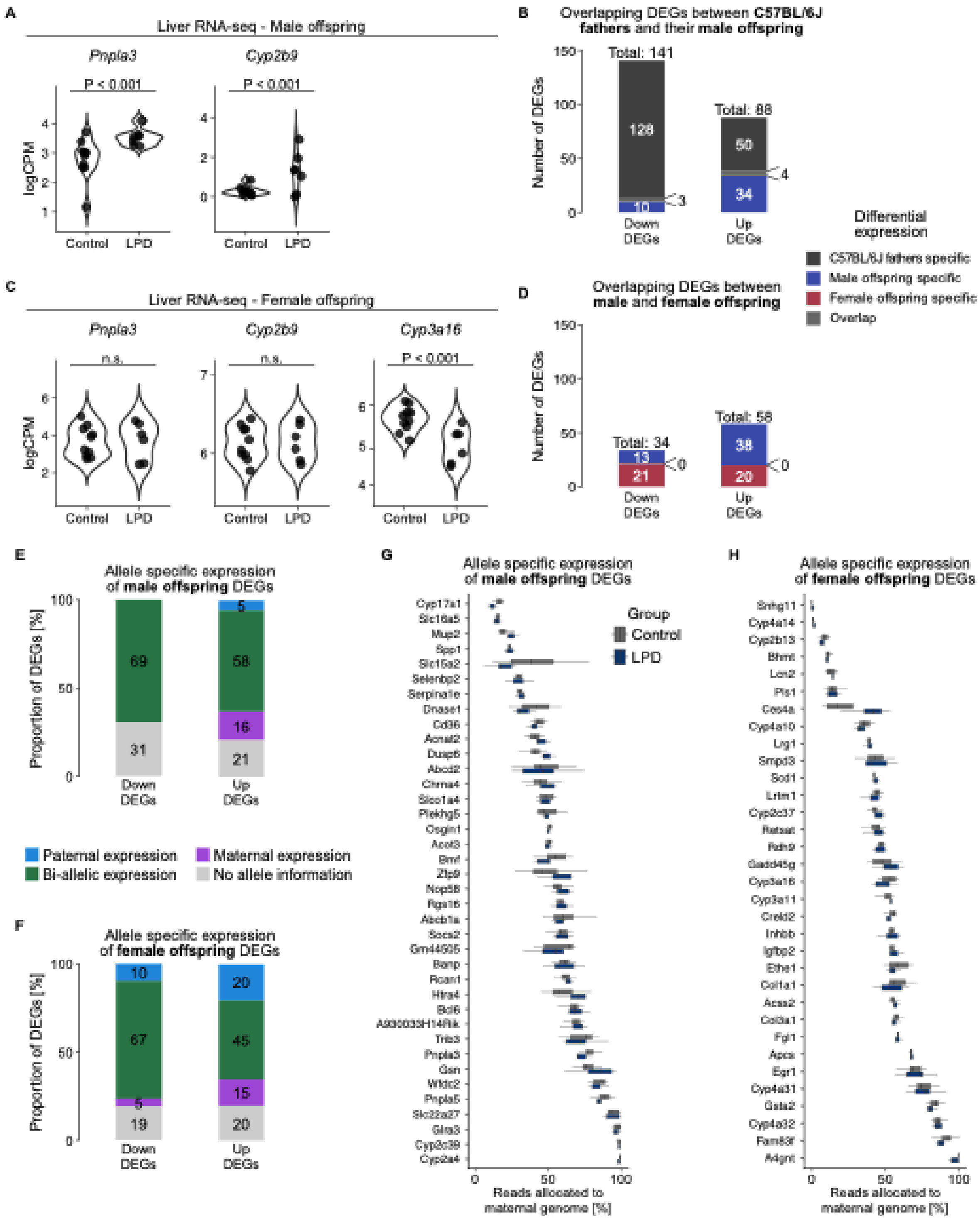

A
Liver RNA-seq - Male offspring
Pnpla3
Cyp2b9
P < 0.001
P < 0.001
logCPM
Control
LPD
B
Overlapping DEGs between C57BL/6J fathers and their male offspring
Number of DEGs
Total: 141
Total: 88
128
10
3
50
34
4
Down DEGs
Up DEGs
Differential expression
C57BL/6J fathers specific
Male offspring specific
Female offspring specific
Overlap
C
Liver RNA-seq - Female offspring
Pnpla3
Cyp2b9
Cyp3a16
n.s.
n.s.
P < 0.001
D
Overlapping DEGs between male and female offspring
Total: 34
Total: 58
13
21
0
38
20
0
E
Allele specific expression of male offspring DEGs
Proportion of DEGs [%]
69
31
5
58
16
21
Paternal expression
Maternal expression
Bi-allelic expression
No allele information
F
Allele specific expression of female offspring DEGs
10
67
5
19
20
45
15
20
G
Allele specific expression of male offspring DEGs
Group
Control
LPD
Cyp17a1
Slc16a5
Mup2
Spp1
Slc15a2
Selenbp2
Serpina1e
Dnase1
Cd36
Acnat2
Dusp6
Abcd2
Chrna4
Slco1a4
Plekhg5
Osgin1
Acot3
Bmf
Zfp9
Nop58
Rgs16
Abcb1a
Socs2
Gm44505
Banp
Rcan1
Htra4
Bcl6
A930033H14Rik
Trib3
Pnpla3
Gsn
Wfdc2
Pnpla5
Slc22a27
Glra3
Cyp2c39
Cyp2a4
Reads allocated to maternal genome [%]
H
Allele specific expression of female offspring DEGs
Snhg11
Cyp4a14
Cyp2b13
Bhmt
Lcn2
Pls1
Ces4a
Cyp4a10
Lrg1
Smpd3
Sod1
Lrtm1
Cyp2c37
Retsat
Rdh9
Gadd45g
Cyp3a16
Cyp3a11
Creld2
Inhbb
Igfbp2
Ethe1
Col1a1
Acss2
Col3a1
Fgl1
Apcs
Egr1
Cyp4a31
Gsta2
Cyp4a32
Fam83f
A4gnt

**Figure S3. Supplementary analyses of gene expression, overlap analyses, and allele-specific expression in hybrid offspring of C57BL/6J males fed LPD.** (**A**) Violin plots showing expression of selected genes in control and LPD groups in male offspring. *P* values are derived from differential expression analyses. (**B**) Stacked bar plots showing the overlap of down- and upregulated DEGs between C57BL/6J males and their male offspring. (**C**) Violin plots showing expression of selected genes in control and LPD groups in female offspring. *P* values are derived from differential expression analyses. (**D**) Stacked bar plots showing the overlap of down- and upregulated DEGs between male and female offspring. (**E**) Stacked bar plots showing the distribution of allele-specific expression among down- and upregulated DEGs in male offspring. (**F**) Boxplots showing the proportion of reads allocated to the maternal genome per sample in control and LPD groups for DEGs in male offspring. Each box represents the distribution of maternal allele expression across samples of a group for a given gene. No gene showed a significant difference in allele-specific expression between groups. (**G**) Stacked bar plots showing the distribution of allele-specific expression among down- and upregulated DEGs in female offspring. (**H**) Boxplots showing the proportion of reads allocated to the maternal genome per sample in control and LPD groups for DEGs in female offspring. Each box represents the distribution of maternal allele expression across samples of a group for a given gene. No gene showed a significant difference in allele-specific expression between groups.

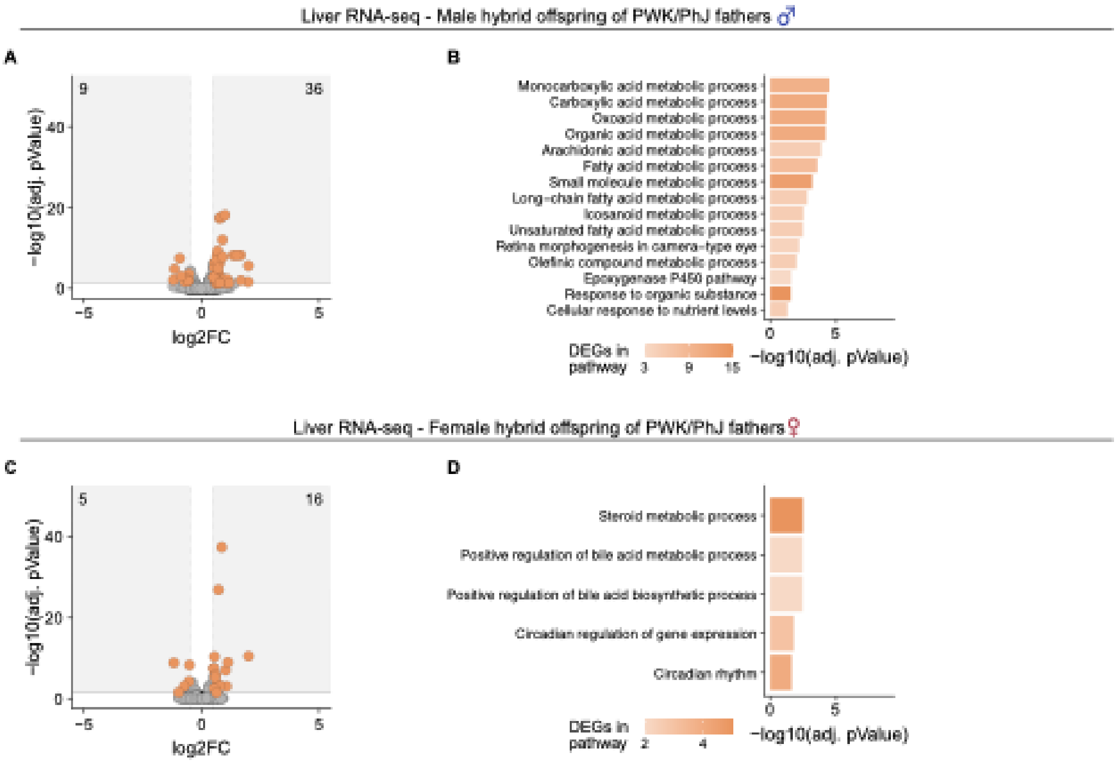


**Figure S4. Analyses of liver transcriptome in offspring of PWK/PhJ males fed LPD.** (**A** and **B**) Effects of paternal LPD on the liver transcriptome of male offspring of PWK/PhJ males. Control: n = 10, LPD: n = 10. (**A**) Volcano plot of DEGs (adjusted *P* < 0.05 and abs(log2FC) > 0.5) in liver of male offspring of PWK/PhJ fathers fed LPD compared to controls. Numbers indicate the count of down- and upregulated DEGs. (**B**) Top 15 biological process GO terms most significantly enriched among DEGs in male offspring. (**C** and **D**) Effects of paternal LPD on the liver transcriptome of female offspring of PWK/PhJ males. Control: n = 10, LPD: n = 11. (**C**) Volcano plot of DEGs (adjusted *P* < 0.05 and abs(log2FC) > 0.5) in liver of female offspring of PWK/PhJ fathers fed LPD compared to controls. Numbers indicate the count of down- and upregulated DEGs. (**D**) Biological process GO terms significantly enriched among DEGs in female offspring.

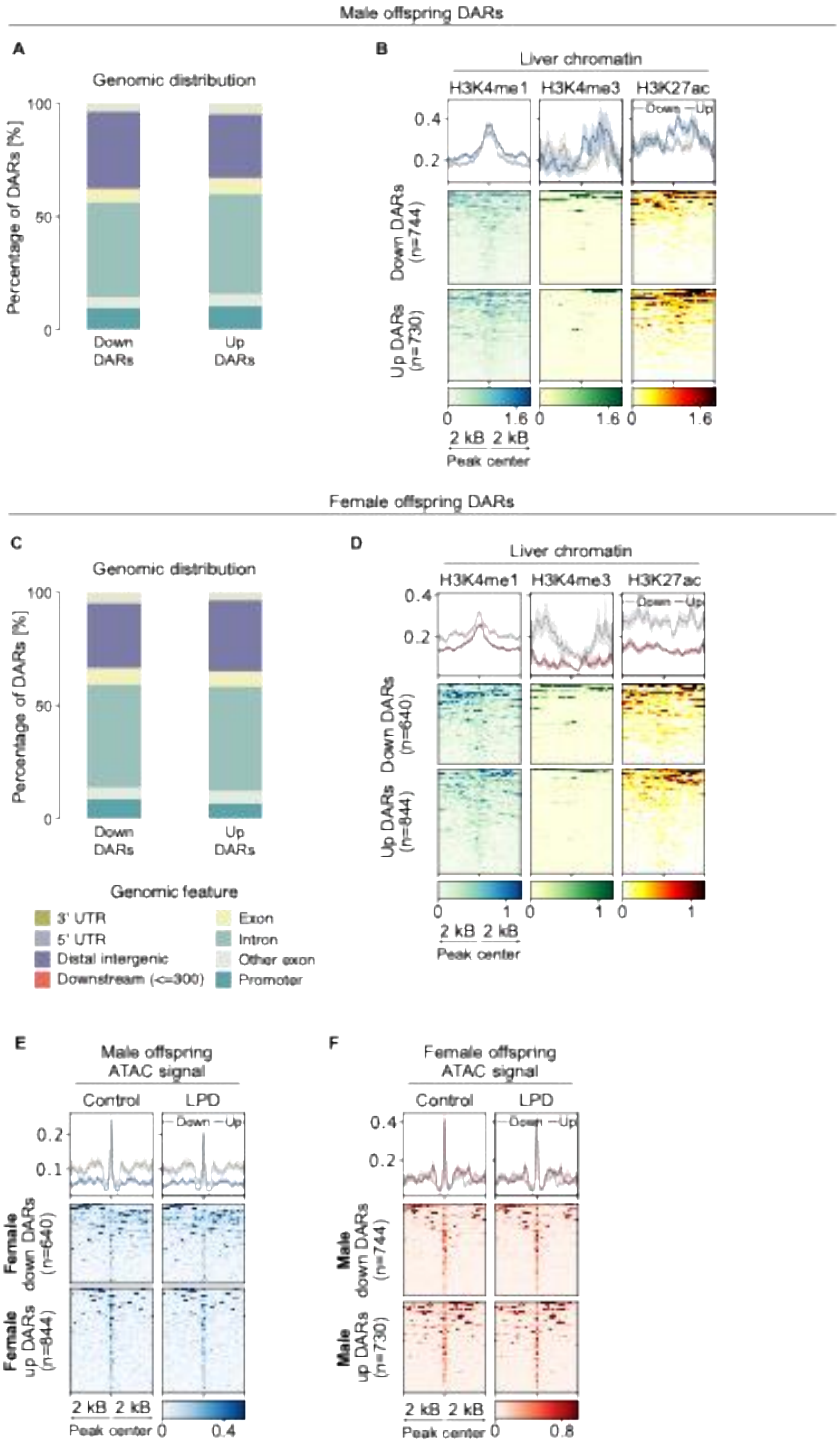

Male offspring DARs
A
Genomic distribution
Percentage of DARs [%]
100
50
0
Down DARs
Up DARs
B
Liver chromatin
H3K4me1
H3K4me3
H3K27ac
Down
Up
0.4
0.2
Down DARs (n=744)
Up DARs (n=730)
0
1.6
2 kB
2 kB
Peak center
Female offspring DARs
C
Genomic distribution
Percentage of DARs [%]
Down DARs
Up DARs
Genomic feature
3' UTR
5' UTR
Distal intergenic
Downstream (<=300)
Exon
Intron
Other exon
Promoter
D
Liver chromatin
Down DARs (n=640)
Up DARs (n=844)
1
E
Male offspring ATAC signal
Control
LPD
Female down DARs (n=640)
Female up DARs (n=844)
0.4
F
Female offspring ATAC signal
Male down DARs (n=744)
Male up DARs (n=730)
0.8

**Figure S5. Supplementary analyses of chromatin accessibility in offspring of C57BL/6J fathers fed LPD.** (**A** and **B**) Genomic annotation of DARs identified in male offspring. (**A**) Stacked bar plot showing the genomic distribution of DARs. (**B**) Heatmaps showing normalized H3K4me1, H3K4me3, and H3K27ac ChIP-seq signal in liver. Each row represents a 4-kb region centered on the DAR midpoint (±2 kb). Public data are from Yue *et al.* (2014) [101]. (**C** and **D**) Genomic annotation of DARs identified in female offspring. (**C**) Stacked bar plot showing the genomic distribution of DARs. (**D**) Heatmaps showing normalized H3K4me1, H3K4me3, and H3K27ac ChIP-seq signal in liver. Each row represents a 4-kb region centered on the DAR midpoint (±2 kb). (**E** and **F**) Heatmaps showing normalized ATAC-seq signal between control and LPD groups in (**E**) female DARs using male offspring ATAC-seq data and (**F**) male DARs using female offspring ATAC-seq data ($P < 0.05$ and abs(log2FC) > 1). Each row represents a 4-kb region centered on the DAR midpoint (±2 kb), ordered by mean chromatin accessibility.

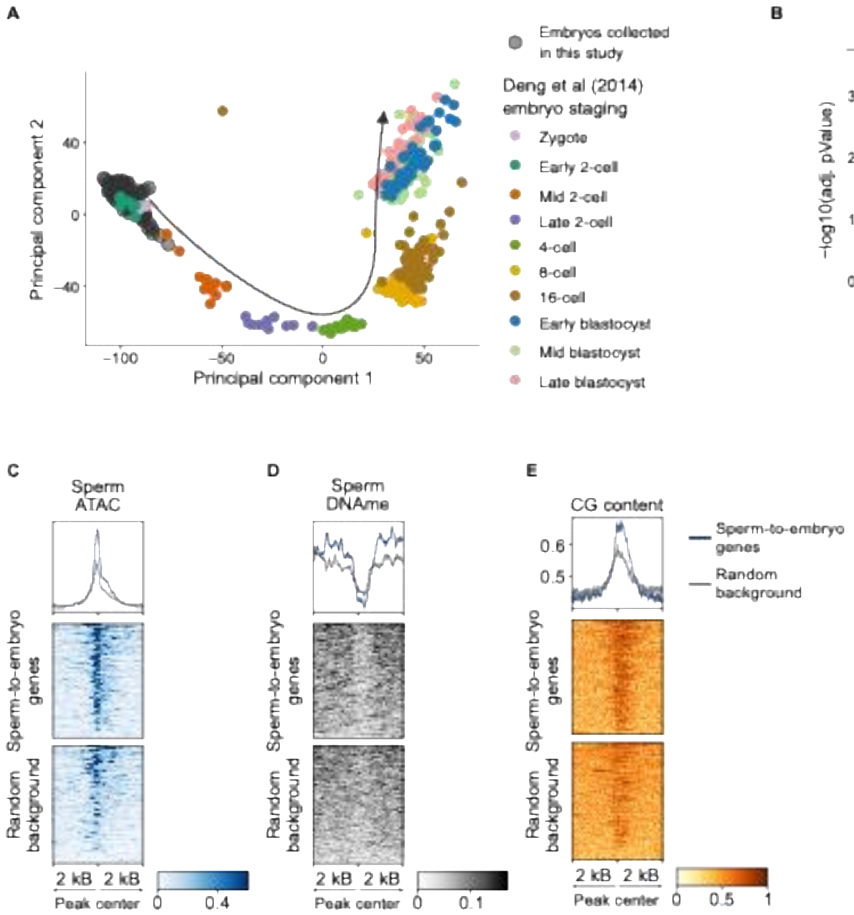
A
B
C
D
E
Embryos collected in this study
Deng et al (2014) embryo staging
Zygote
Early 2-cell
Mid 2-cell
Late 2-cell
4-cell
8-cell
16-cell
Early blastocyst
Mid blastocyst
Late blastocyst
Principal component 1
Principal component 2
Sperm long RNA-seq
−log10(adj. pValue)
log2FC
Sperm ATAC
Sperm DNAme
CG content
Sperm-to-embryo genes
Random background
2 kB
Peak center

**Figure S6. Supplementary analyses of early two-cell embryos, sperm transcriptome, and chromatin features.** (**A**) Principal component analysis (PCA) plot showing staging of preimplantation embryos, with embryos collected in this study mapped onto the reference dataset. Public data are from Deng *et al.* (2014) [43]. (**B**) Volcano plot of DEGs (adjusted $P < 0.05$ and abs(log2FC) > 0.5) in sperm from males fed LPD compared to controls. Numbers indicate the count of down- (*Rhox3e*, *Muc21*, *Chd8*) and upregulated (*Flt3l*) DEGs. Control: n = 8, LPD: n = 8. (**C**) Heatmap showing normalized ATAC-seq signal in sperm. Each row represents a 4-kb region centered on the TSS of sperm-to-embryo or random background genes (±2 kb), ordered by mean ATAC-seq signal. Background genes were randomly chosen from those expressed in early two-cell embryos. (**D**) Heatmap showing normalized DNAme signal in sperm. Each row represents a 4-kb region centered on the TSS of sperm-to-embryo or random background genes (±2 kb), ordered by mean DNAme signal. Background genes were randomly chosen from those expressed in early two-cell embryos. (**E**) Heatmap showing CG content. Each row represents a 4-kb region centered on the TSS of sperm-to-embryo or random background genes (±2 kb), ordered by mean CG content. Background genes were randomly chosen from those expressed in early two-cell embryos.

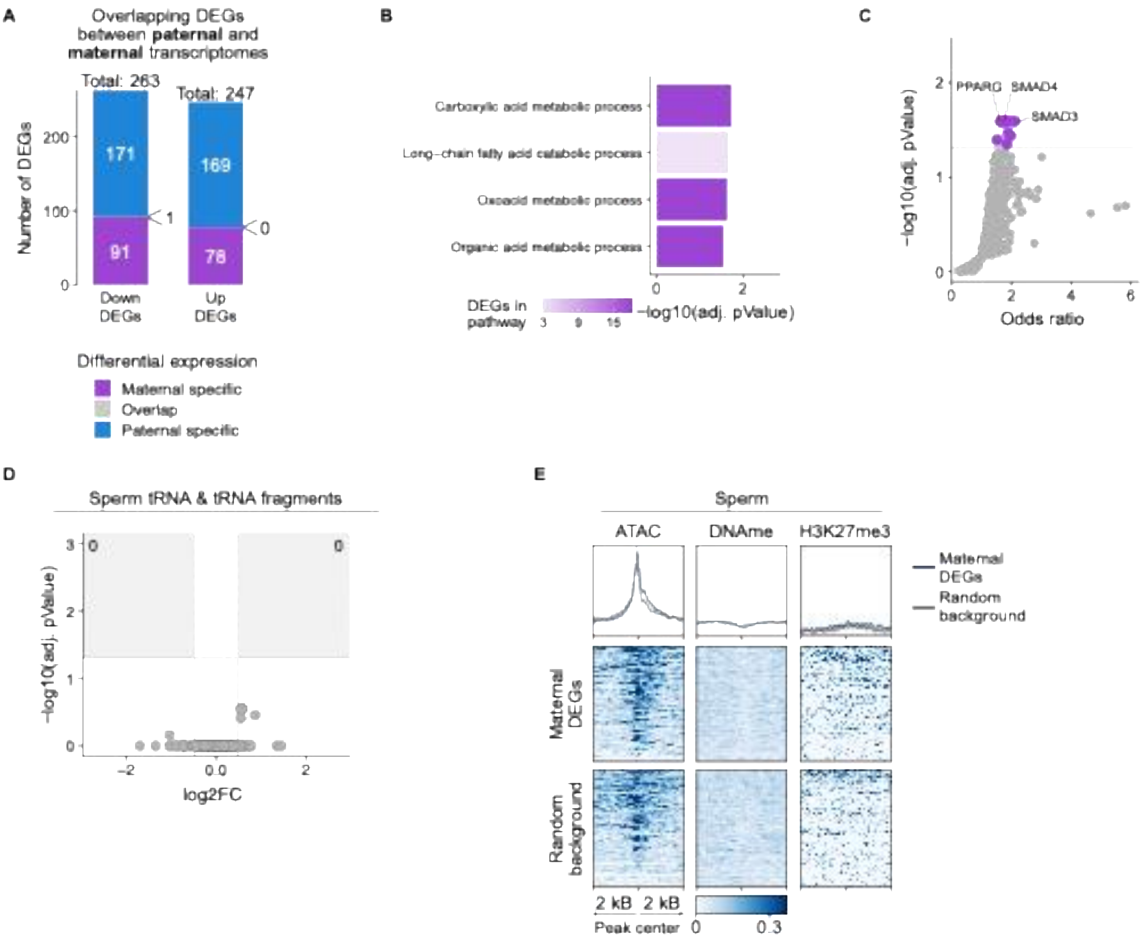


**Figure S7. Analyses of DEGs in the maternal transcriptome of early two-cell embryos and sperm small RNAs.** (**A**) Stacked bar plots showing the overlap of down- and upregulated DEGs between paternal and maternal transcriptomes of early two-cell embryos. Overlapping downregulated gene: *Zfp748* (**B**) Biological process GO terms significantly enriched among DEGs in the maternal transcriptome of early two-cell embryos. (**C**) TFs with targets enriched among DEGs in the maternal transcriptome of early two-cell embryos. Colored points indicate significantly enriched TFs after multiple testing correction. (**D**) Volcano plot of differentially expressed tRNAs (adjusted $P < 0.05$ and abs(log2FC) > 0.5) in sperm from fathers fed LPD compared to controls. Numbers indicate the count of down- and upregulated tRNAs. Control: n = 9, LPD: n = 9. (**E**) Heatmaps showing normalized ATAC-seq, DNAme, and H3K27me3 ChIP-seq signal in sperm. Each row represents a 4-kb region centered on the TSS of maternal DEGs or random background genes (±2 kb), ordered by mean ChIP-seq signal. Background genes were randomly chosen from those expressed in early two-cell embryos. H3K27me3 public data are from Zheng *et al.* (2016) [60].

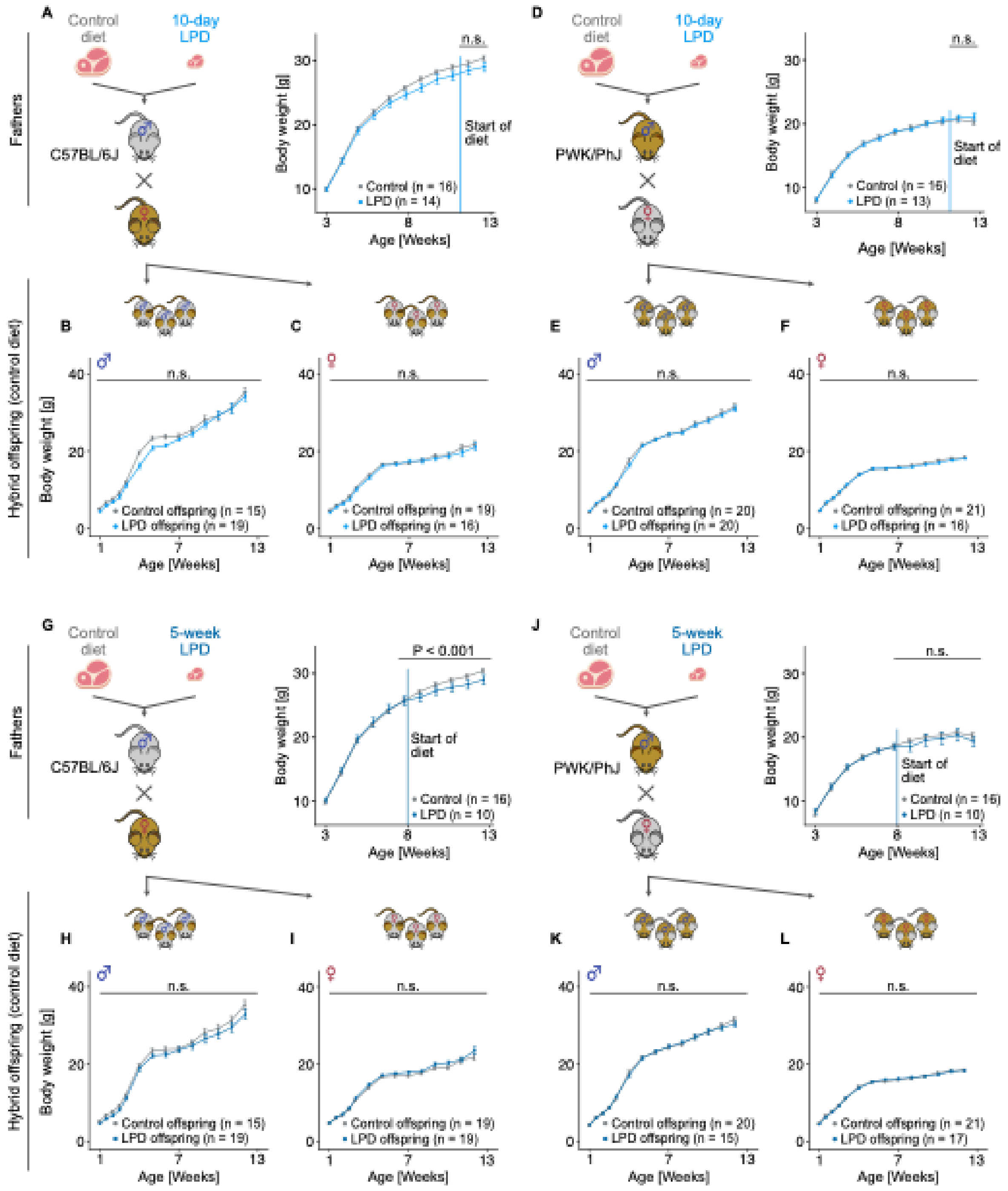

A
Control diet
10-day LPD
Fathers
C57BL/6J
n.s.
Start of diet
Body weight [g]
Control (n = 16)
LPD (n = 14)
Age [Weeks]
D
Control diet
10-day LPD
PWK/PhJ
n.s.
Start of diet
Control (n = 16)
LPD (n = 13)
B
C
E
F
Hybrid offspring (control diet)
n.s.
Control offspring (n = 15)
LPD offspring (n = 19)
Control offspring (n = 19)
LPD offspring (n = 16)
Control offspring (n = 20)
LPD offspring (n = 20)
Control offspring (n = 21)
LPD offspring (n = 16)
G
Control diet
5-week LPD
C57BL/6J
P < 0.001
Start of diet
Control (n = 16)
LPD (n = 10)
J
Control diet
5-week LPD
PWK/PhJ
n.s.
Start of diet
Control (n = 16)
LPD (n = 10)
H
I
K
L
Control offspring (n = 15)
LPD offspring (n = 19)
Control offspring (n = 19)
LPD offspring (n = 19)
Control offspring (n = 20)
LPD offspring (n = 15)
Control offspring (n = 21)
LPD offspring (n = 17)

**Figure S8. Analyses of 10-day and 5-week LPD effects on body weight growth in C57BL/6J and PWK/PhJ males and their offspring.** (**A**–**C**) Experimental design and body weight trajectories for (**A**) C57BL/6J fathers fed a control diet or a 10-day LPD, and their (**B**) male and (**C**) female hybrid offspring. (**D**–**F**) Experimental design and body weight trajectories for (**D**) PWK/PhJ fathers fed a control diet or a 10-day LPD, and their (**E**) male and (**F**) female hybrid offspring. (**G**–**I**) Experimental design and body weight trajectories for (**G**) C57BL/6J fathers fed a control diet or a 5-week LPD, and their (**H**) male and (**I**) female hybrid offspring. (**J**–**L**) Experimental design and body weight trajectories for (**J**) PWK/PhJ fathers fed a control diet or a 5-week LPD, and their (**K**) male and (**L**) female hybrid offspring. Data are presented as mean ± SEM. Statistical analyses for exposed males used a mixed-effects model (*Bodyweight* ~ Group:Age + Age + (1|Animal) + (1|Litter)) and included only data from the period of dietary exposure. Statistical analyses for hybrid offspring used a mixed-effects model (*Bodyweight* ~ Group:Age + Age + (1|Animal) + (1|Father)). All *P* values correspond to the Group:Age interaction, representing changes in body weight during dietary exposure (fathers) or during development (offspring).

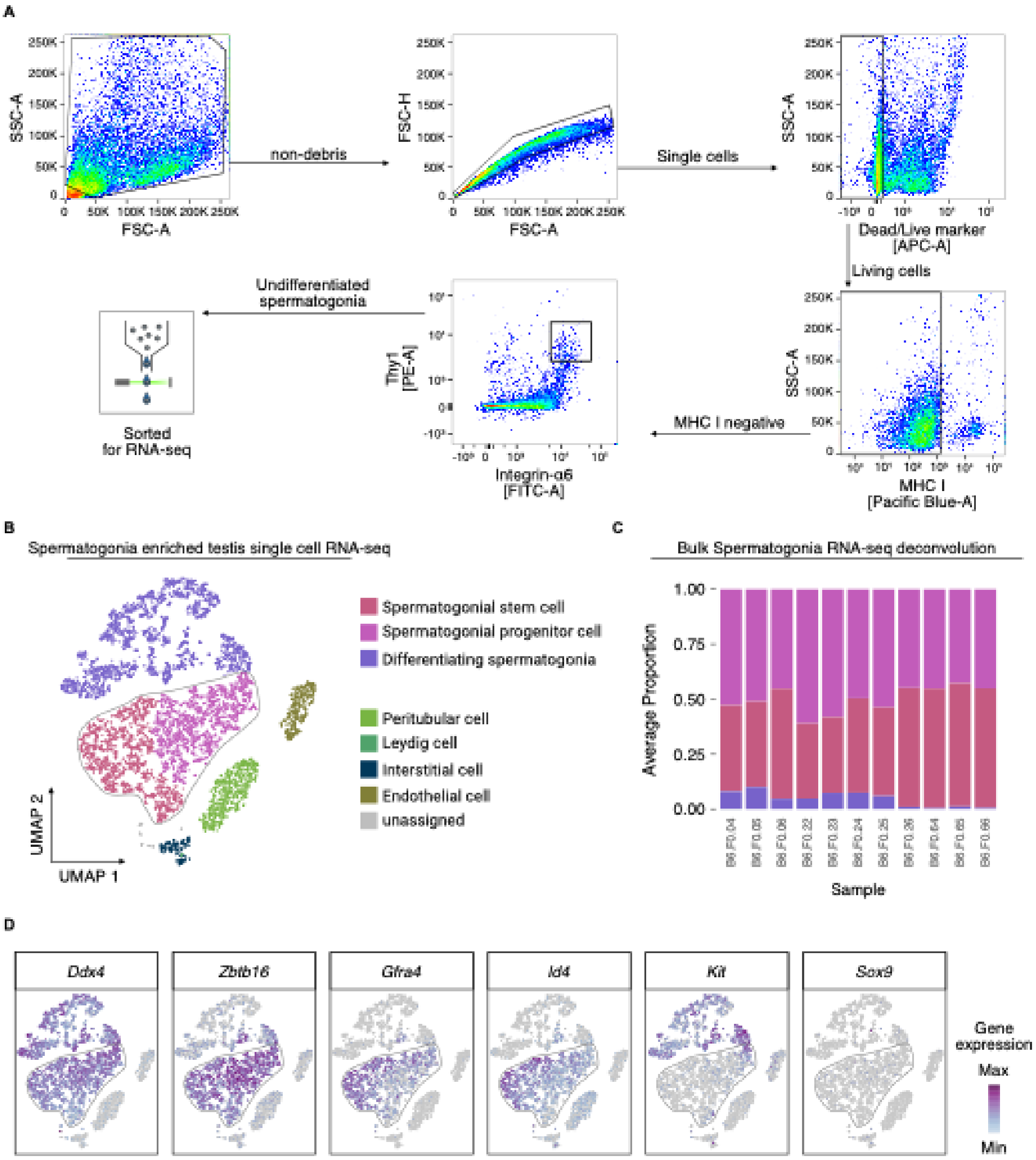

A
SSC-A
FSC-A
non-debris
FSC-H
FSC-A
Single cells
SSC-A
Dead/Live marker
[APC-A]
Living cells
SSC-A
MHC I
[Pacific Blue-A]
MHC I negative
Thy1
[PE-A]
Integrin-α6
[FITC-A]
Undifferentiated
spermatogonia
Sorted
for RNA-seq
B
Spermatogonia enriched testis single cell RNA-seq
Spermatogonial stem cell
Spermatogonial progenitor cell
Differentiating spermatogonia
Peritubular cell
Leydig cell
Interstitial cell
Endothelial cell
unassigned
UMAP 2
UMAP 1
C
Bulk Spermatogonia RNA-seq deconvolution
Average Proportion
1.00
0.75
0.50
0.25
0.00
Sample
D
Ddx4
Zbtb16
Gfra4
Id4
Kit
Sox9
Gene
expression
Max
Min

**Figure S9. Flow cytometry sorting and deconvolution-based characterization of undifferentiated spermatogonia for RNA-seq.** (**A**) Gating strategy used to sort live MHC I–negative, α6-integrin–positive, and Thy1–positive testicular cells to enrich for undifferentiated spermatogonia for RNA-seq. (**B**) UMAP of a testicular single-cell RNA-seq dataset enriched for spermatogonia, with annotated cell types. Public data are from Hermann *et al.* (2018) [57]. (**C**) Bar plot showing estimated cellular proportions for all sorted bulk RNA-seq libraries. Cellular composition was estimated using deconvolution analysis with the single-cell RNA-seq dataset shown in (**B**) as reference. (**D**) Feature plots showing expression of marker genes for early undifferentiated spermatogonia, differentiating spermatogonia, and somatic cell populations.

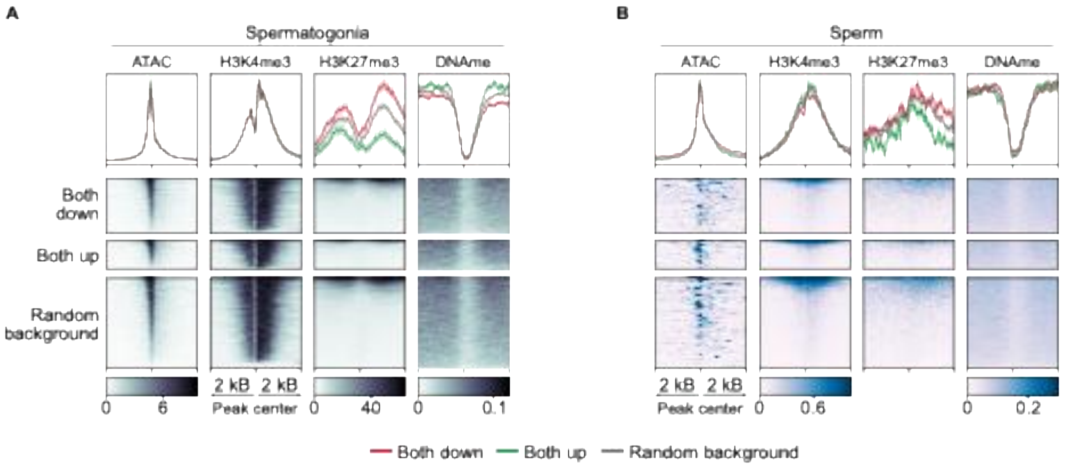


**Figure S10. Chromatin modifications at same-directional genes in undifferentiated spermatogonia and sperm.** (**A** and **B**) Heatmaps showing normalized ATAC-seq, H3K4me3 and H3K27me3 ChIP-seq, and DNAme signals in (**A**) spermatogonia and (**B**) sperm. Each row represents a 4-kb region centered on the TSS of same-directional genes or random background genes (±2 kb), ordered by mean signal. Background genes were randomly chosen from those expressed in spermatogonia. Public data are from Lazar-Contes *et al.* (2025) [56] (spermatogonia ATAC-seq), Cheng *et al.* (2020) [100] (spermatogonia H3K4me3 and H3K27me3 from 6-days old mice), Zoch *et al.* (2020) [102] (spermatogonia DNAme), Zhang *et al.* (2016) [99] (sperm H3K4me3), and Zheng *et al.* (2016) [60] (sperm H3K27me3).
